# Self-assembling shell proteins PduA and PduJ have essential and redundant roles in bacterial microcompartment assembly

**DOI:** 10.1101/2020.07.06.187062

**Authors:** Nolan W. Kennedy, Svetlana P. Ikonomova, Marilyn Slininger Lee, Henry W. Raeder, Danielle Tullman-Ercek

## Abstract

Protein self-assembly is a common and essential biological phenomenon, and bacterial microcompartments present a promising model system to study this process. Bacterial microcompartments are large, protein-based organelles which natively carry out processes important for carbon fixation in cyanobacteria and the survival of enteric bacteria. These structures are increasingly popular with biological engineers due to their potential utility as nanobioreactors or drug delivery vehicles. However, the limited understanding of the assembly mechanism of these bacterial microcompartments hinders efforts to repurpose them for non-native functions. Here, we comprehensively investigate proteins involved in the assembly of the 1,2-propanediol utilization bacterial microcompartment from *Salmonella enterica* serovar Typhimurium LT2, one of the most widely studied microcompartment systems. We first demonstrate that two shell proteins, PduA and PduJ, have a high propensity for self-assembly upon overexpression, and we provide a novel method for self-assembly quantification. Using genomic knock-outs and knock-ins, we systematically show that these two proteins play an essential and redundant role in bacterial microcompartment assembly that cannot be compensated by other shell proteins. At least one of the two proteins PduA and PduJ must be present for the bacterial microcompartment shell to assemble. We also demonstrate that assembly-deficient variants of these proteins are unable to rescue microcompartment formation, highlighting the importance of this assembly property. Our work provides insight into the assembly mechanism of these bacterial organelles and will aid downstream engineering efforts.

## Introduction

Proteins play integral and myriad roles in biological functions. Their study spans and interconnects a vast number of research fields, including genome editing with CRISPR and enzyme-driven bioremediation. Just as proteins tie together research fields, proteins themselves interact and work together to build intricate assemblies and execute complex functions. Bacterial microcompartments (MCPs) are one example of protein assembly contributing to a cellular function. These are large protein-based organelles found in as many as two thirds of all bacterial phyla [1–3]. There are multiple subtypes of MCPs, including carboxysomes, which are used for carbon fixation, and numerous types of metabolic MCPs used for breaking down unique carbon sources for energy. Metabolic MCPs are often found in enteric bacteria, including harmless microbes present in the human gut microbiome as well as pathogens such as *Salmonella enterica* and *Escherichia coli.* These organelles encapsulate enzymes necessary to metabolize niche carbon sources found in the gut of hosts [4,5]. The encapsulated enzymes often produce a toxic intermediate, which is thought to be retained within the assembled protein shell [6,7]. Thus, MCPs protect the cell from harmful levels of toxic intermediates and facilitate safe, efficient metabolism.

One of the most well-studied MCPs is the 1,2-propanediol utilization (Pdu) MCP from *Salmonella enterica* serovar Typhimurium LT2. A single 22-gene operon encodes the MCP, which includes enzymes for 1,2-propanediol (1,2-PD) metabolism and cofactor recycling, as well as structural proteins that comprise the MCP shell [4,8–10]. The Pdu MCP is an irregular polyhedral structure (Figure 1), which has a maximum diameter that ranges from approximately 75-200 nm [11]. While MCPs are believed to play a role in the survival and proliferation of *Salmonella,* these structures have also grown in popularity due to their potential utility in metabolic engineering [12–14]. Synthetic biologists are repurposing Pdu MCPs to encapsulate non-native, industrially relevant pathways to increase pathway flux [12]. MCPs offer benefits such as a known, modular mechanism for enzyme encapsulation and the native ability to recycle cofactors while preventing off-target interactions in the cytosol [9,10,15–18]. For researchers to fully harness these benefits and capabilities of MCPs, knowledge of MCP biogenesis is critical, but pieces are still missing to this mechanistic puzzle.

**Figure 1.**
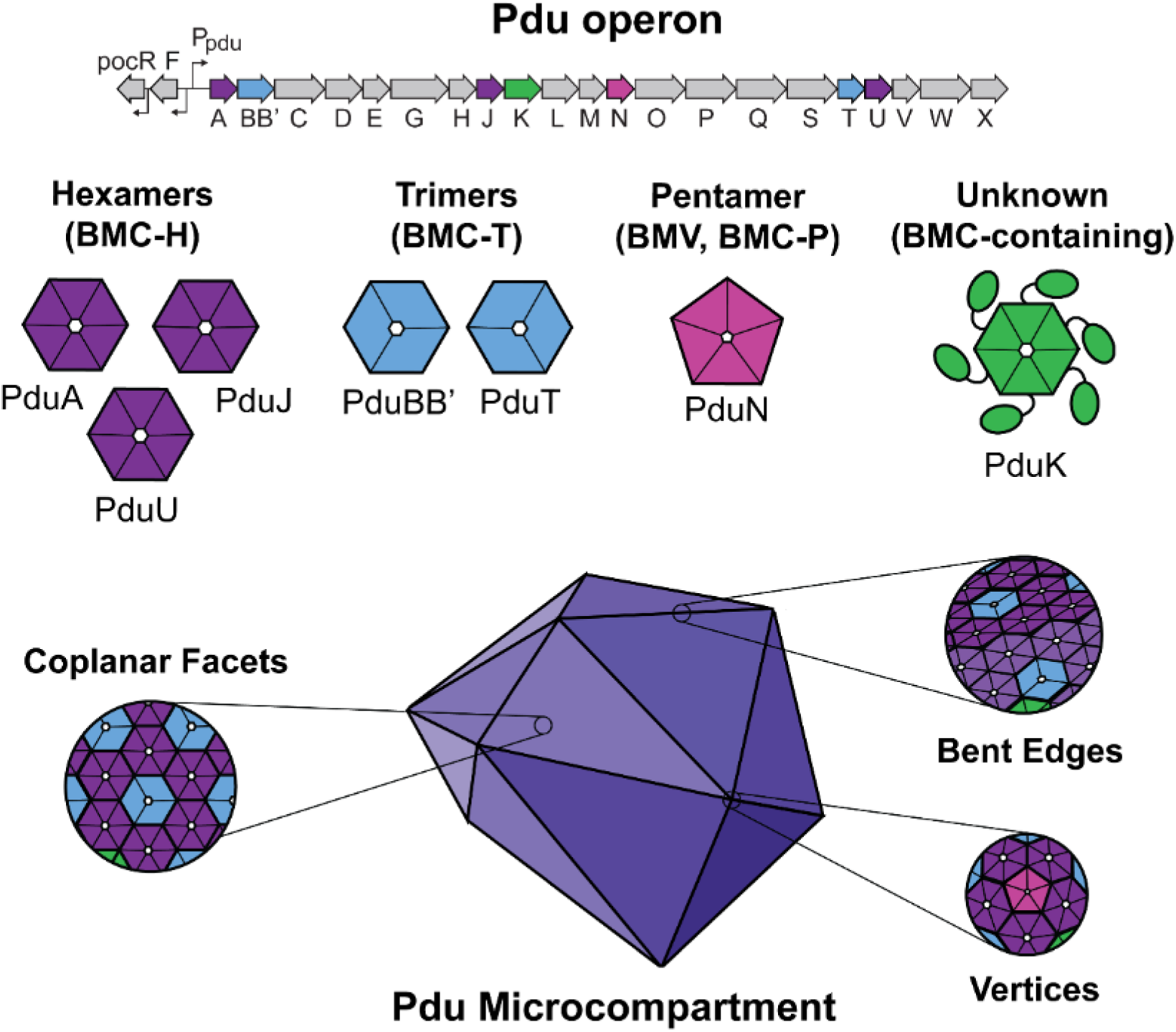
Different shell proteins make up the Pdu MCP shell. Schematic representation of the eight different genomically-encoded shell proteins within the Pdu operon. These proteins have four different basic architectures, including hexamers (BMC-H), trimers (BMC-T), pentamers (BMV, BMC-P), or unknown. Together, these assemble into a mosaic shell which contains flat facets, bent edges, and vertices.

An area in need of investigation is the mechanism of Pdu MCP shell assembly. The *pdu* operon contains open reading frames for at least eight shell proteins (Figure 1). These shell proteins come in three basic architectures: hexamers, trimers, and pentamers. PduA, PduJ, and PduU are experimentally verified to contain a bacterial microcompartment (BMC/ pfam00936)-domain; six copies of each protein (BMC-H) form hexamers based on crystal structures [19–21]. These form flat, six-sided hexagonal disks that tile into the facets of the full MCP shell (Figure 1). PduA and PduJ are two of the most abundant members of the Pdu MCP shell [8,22], and have been predicted to play a variety of roles, such as MCP assembly [23], control of substrate diffusion through their central open pores [24,25], and enzyme encapsulation [26,27]. PduU is a minor shell component and its role within the Pdu MCP is still unknown, though it is theorized to not play role with substrate diffusion as its central pore is capped by a β-barrel [21]. These BMC-H proteins are predicted to play a major role in MCP assembly based on structural studies of a related MCP system from *Haliangium ochraceum,* which showed that they interact with the other MCP shell proteins and can adopt both a coplanar and bent interaction orientation (Figure 1) [28]. PduBB’ and PduT are experimentally verified as trimers (BMC-T) (Figure 1) [19,29]. These trimeric proteins contain two tandem BMC-domains that are physically linked in a single, contiguous polypeptide chain. PduBB’ is one of the most abundant members of the Pdu MCP shell (after PduA and PduJ) [8,22], and is predicted to play a role in regulating substrate diffusion (possibly with a gated pore mechanism [30,31]), as well as in enzyme encapsulation [32]. The role of PduT is less well-understood, but it has been hypothesized to use a central iron-sulfur cluster for electron transport across the MCP shell [29]. PduN encodes a single bacterial microcompartment vertex domain (BMV-domain, Pfam03319), and is a minor shell component [8,22]. While a crystal structure has not been solved of PduN, other BMV-domain proteins form pentameric (BMC-P) structures that are thought to occupy the vertices of the irregular polyhedral MCP shells [30] similar to carboxysomes (Figure 1). Interestingly, homologous BMV-domain proteins have been crystalized as pentamers or hexamers, indicating a potential dual function within the shell [30]. Finally, PduK is a putative shell protein that contains a single BMC-domain as well as a long C-terminal extension [1]. While the function and architecture of PduK is still not known, it is putatively a hexameric shell protein and potentially plays a role in enzyme encapsulation [12]. While some evidence exists for a role for each of the Pdu MCP shell proteins, how these proteins work together to assemble into a final closed, polyhedral structure is not yet known. For example, some BMC-H shell proteins such as PduA and PduJ can form various extended tubes or filaments when overexpressed [35–37] and are likely a propagation of the natural bent interaction between hexamers at the interface of two MCP shell facets (Figure 1) [28]. Current models based on related structures, such as carboxysomes, suggest that the shell assembles around a preformed core (or concomitantly with the core) [33,34], but which proteins are responsible for buildup of the shell, or how shell assembly might be initiated, is not known for the Pdu or other systems.

In this work, we confirm that two of the Pdu shell proteins, PduA and PduJ, have a unique, intrinsic propensity for self-assembly into long tubes upon overexpression. We hypothesized that the tendency to self-assemble makes PduA and PduJ important for MCP assembly as a whole, and that they may be the components necessary to initiate shell assembly. To test this hypothesis, we developed a microscopy-based method for the rapid quantification of this self-assembly property that does not require protein purification. We then performed genomic knockouts of *pduA* and *pduJ* and demonstrate that these two genes are essential and redundant for MCP assembly. We further show that the loss of assembly cannot be overcome by other Pdu shell proteins, and that assembly-deficient mutants of these proteins do not recover MCP assembly. Findings from this study also provide a new, non-assembled MCP control for studies investigating or modifying the MCP shell or evaluating the encapsulation of heterologous pathways. Overall, this work determined essential components of MCP shell formation, provides insight into the assembly process, and offers new techniques to use when engineering MCPs.

## Results

### PduA and PduJ self-assemble into tubes when overexpressed

The *pdu* operon in *Salmonella enterica* serovar Typhimurium LT2 contains open reading frames for eight shell proteins *(pduA, -B, -B’, -J, -K, -N, -T,* and *-U)* [4,8,38]. Some of these shell proteins self-assemble into large, multimeric structures when overexpressed [23,35–37,39,40]. Indeed, when we overexpress the Pdu shell proteins PduA and PduJ in *E. coli* BL21 cells, we observe that they self-assemble into long, hollow tubes (Figure 1A, 2B). Such elongated assemblies are predicted by theory and are also observed in the virus world. These tubes are observable in the cytoplasm when cells overexpressing PduA or PduJ are sectioned, stained, and imaged (Figure 2A, 2B). In cell sections, PduA and PduJ tubes appear as either hollow circles (transverse section), or as long, parallel stripes (longitudinal sections), which likely correspond to the walls of the protein tubes (Figure 2A, 2B). We observed that these PduA and PduJ tubes packed tightly together into bundles within the *E. coli* cytoplasm. These bundles contained an average of 17 and 10 tubes for PduA and PduJ, respectively, and ranged from 4 to 60 tubes per bundle. Quantification of the tube diameters showed that PduA and PduJ tubes are 19.5 ± 2.5 nm and 20.4 ±5.1 nm in diameter, respectively. These results correspond well to previous characterizations of PduA tubes and the tubes of a PduA homolog, RmmH, all of which reported approximate diameters of 20 nm [35,39,41]. These reports suggested either a parallel or alternating arrangement of protein hexamers, with 8-12 units per turn. Given the size of PduA hexamers (~5 nm diameter), and observed tube diameter (~20 nm), we propose 12 units per turn with an external angle of 30 degrees. The structure of a homologous, simplified MCP exhibits a 30 degree angle between hexamers at the bent interface (Figure 1), leading us to hypothesize that PduA hexamers are adopting this conformation in the tubes. [28]. PduA and PduJ tubes are also visible via transmission electron microscopy (TEM) after purification from cells using differential centrifugation (Figure 2B).

**Figure 2.**
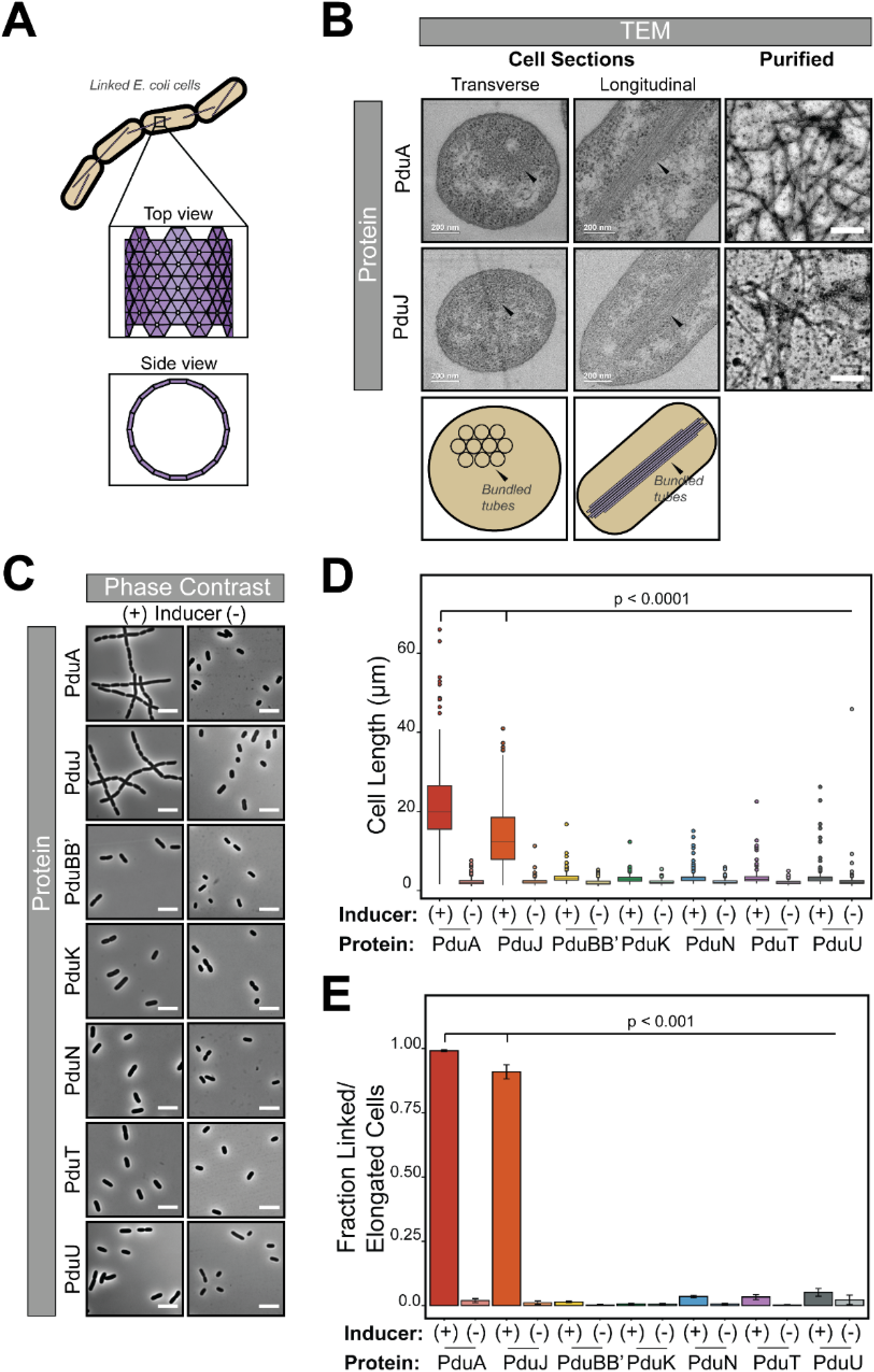
PduA and PduJ self-assemble into tubes when overexpressed. (A) Schematic representation of PduA or PduJ tubes in *E. coli* cells. These appear as long structures which run along the long axis of the cell body (top view), or as hollow circles when sectioned down the center (side view). (B) TEM of PduA and PduJ self-assembled into long tubes. Cell sections of *E. coli* cells show PduA (top left and middle) and PduJ (bottom left and middle) self-assembled into tubes when overexpressed. These are visible in both transverse (left) and longitudinal (middle) cell sections (scale bar = 200 nm). Black arrows indicate the tubes. Self-assembled PduA and PduJ tubes can also be purified using differential centrifugation and visualized via TEM (right top and bottom, respectively) (scale bar = 500 nm). (C) Cells induced (+ inducer) or uninduced (-inducer) for overexpressing Pdu shell proteins observed using phase contrast microscopy (scale bar = 5 μm). (D) Length of cells induced (+) or uninduced (-) for overexpression of Pdu shell proteins. Cells overexpressing PduA or PduJ (+ inducer) are significantly longer than cells overexpressing other Pdu shell proteins or not overexpressing shell protein (-inducer) (p < 0.0001, t-test). Measurements were taken from three biological replicates. (E) Fraction of linked or elongated cells in population of cells induced (+) or uninduced (-) for Pdu shell protein overexpression. A significantly higher fraction of cells overexpressing PduA or PduJ (+ inducer) are in linkages of three or more cells or longer than 10 μm (p < 0.001, t-test). Counts were taken from three biological replicates (>80 cells counted per strain per replicate), and error bars represent standard error of the means.

Cells overexpressing PduA or PduJ exhibited a severe cell division defect upon induction and expression (Figure 2C). Cells with this defect appeared primarily as long chains of contiguous cells, but occasionally appeared as highly-elongated single cells (Figure 2C). Cells induced for PduA or PduJ expression were significantly longer than uninduced cells *(i.e.,* no arabinose added to cultures containing *pduA* or *pduJ* open reading frames in arabinose-inducible vectors) (p < 0.0001) (Figure 2D). On average, induced cells were approximately 13-21 μm long, while uninduced cells were approximately 2 μm long (p < 0.0001) (Figure 2D). Over 90% of cells expressing PduA or PduJ existed in chains of three or more cells or single cells greater than 10 μm long, while fewer than 5% of uninduced cells were linked or elongated (p < 0.001) (Figure 2E). This method for assembly quantification via counting linked or elongated cells was more rapid than measuring cell length using an imaging software, and the two methods correlated well (R^2^ = 0.94) (Figure S1A).

To determine whether the cell division defect is unique to PduA and PduJ, we overexpressed the other Pdu shell proteins (Figure 2C-E).While we expected that shell proteins with stronger homology to PduA and PduJ would be most likely to assemble into tubes, we tested all putative shell proteins for assembly, including the BMC-domain-containing PduK and the BMV-domain protein PduN. However, fewer than 6% of all uninduced cells or cells induced for PduB, - K, -N, -T, or -U expression were linked or elongated, significantly fewer than cells expressing PduA or PduJ (p < 0.001) (Figure 2E). Cells expressing shell proteins other than PduA or PduJ were also significantly shorter on average (p < 0.0001) (Figure 2D). We confirmed expression of all FLAG-tagged shell proteins by western blot (Figure S1B), which demonstrated that the observed differences in assembly were not due to lack of expression. Together, in support of previous studies, these results suggest that PduA and PduJ but not the other Pdu shell proteins self-assemble into extraordinarily long tubes that cause a severe cell-division defect when overexpressed in *E. coli* [23,27,35–37,39,40]. While previous works demonstrated that other shell proteins can form smaller-scale aberrant structures upon overexpression [35–37], our results emphasize major differences in the nature and degree of self-assembled PduA and PduJ structures relative to those formed by other shell proteins. This finding led us to hypothesize that PduA- and PduJ-specific self-assembly into tubes is an important property of these proteins, which is linked to overall MCP assembly.

### PduA and PduJ are essential and redundant for microcompartment assembly

We hypothesized that PduA and PduJ are essential for MCP assembly, but previous studies suggest that individual knockouts of either *pduA* or *pduJ* from the *pdu* operon still form MCPs [20,42]. Due to the structural and sequence similarity of these two proteins (Figure S2), we further theorized that PduA and PduJ may play redundant roles in MCP assembly and compensatory effects mask the importance of these proteins in single knockouts. To test these ideas, we created a double knockout strain of LT2 that contained neither *pduA* nor *pduJ* (ΔA ΔJ) (Figure 3A). We were unable to purify MCPs from this strain, as indicated by the absence of the standard wild-type (WT) MCP banding pattern observed via sodium dodecyl sulfate-polyacrylamide gel electrophoresis (SDS-PAGE) (Figure 3B). In contrast, the banding pattern in the single *pduA* (ΔA) and *pduJ* (ΔJ) knockouts is similar to that of the WT MCPs, apart from the missing PduA and PduJ bands at ~10 kDa. TEM on samples purified from the ΔA ΔJ double knockout strain showed only aggregates, which lacked the distinguished, flat, angular boundaries characteristic of WT MCPs (Figure 3C), whereas MCPs purified from the individual knockout strains (ΔA or ΔJ) appeared morphologically similar to WT MCPs (Figure 3C). Note that Pdu MCPs are irregular and polyhedral in nature, and therefore samples are morphologically heterogeneous.

**Figure 3.**
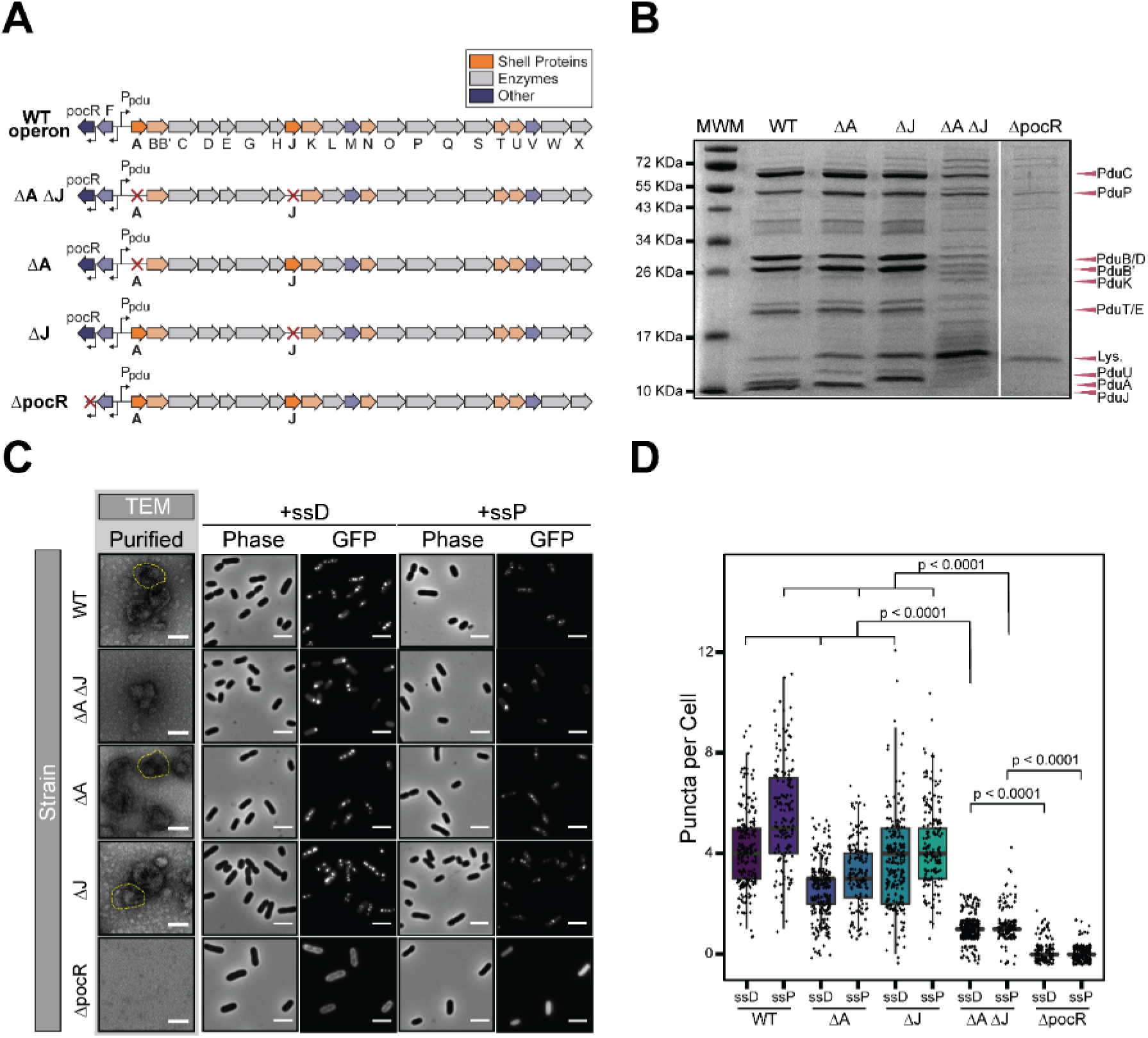
Double knockout of *pduA* and *pduJ* impairs MCP formation. (A) Schematic representation of *pdu* operon and *pdu* gene knockouts. (B) Coomassie-stained SDS-PAGE gel of purified MCPs from different strains with Pdu MCP components labelled. “MWM” is the molecular weight marker and “Lys.” is lysozyme from the lysis buffer. (C) (Left) TEM of purified MCPs from different strains. The image of WT MCPs shows the distinct irregular polyhedral shape and borders we expect from properly formed MCPs. Example MCPs are outlined in yellow. Scale bar = 100 nm. (Right) Phase contrast and GFP fluorescence microscopy images of strains encapsulating either ssD-GFP or ssP-GFP. Row labels indicate the strain and column labels indicate the type of microscopy and the signal sequence appended to GFP. Scale bars = 3 μm. (D) Quantification of puncta per cell for strains encapsulating either ssD-GFP or ssP-GFP. WT, ΔA, and ΔJ strains had significantly greater numbers of puncta per cell than the ΔA ΔJ strain, regardless of signal sequence used to tag GFP (p < 0.0001, t-test). Counts were collected from three biological replicates (>50 cells counted per strain per replicate). Overlaid data points represent the number of puncta for an individual cell and are distributed randomly around the whole number value (y-axis) to facilitate visualization.

To further investigate the role of PduA and PduJ in MCP formation, we tested encapsulation of GFP in MCPs using a microscopy-based encapsulation assay. GFP was directed to MCPs by translationally fusing the protein with native signal sequences from PduD (ssD) and PduP (ssP) [15,16,43]. Strains capable of forming MCPs with encapsulated GFP display bright, fluorescent puncta in the cell cytoplasm (Figure 3C) while cells with uninduced *pdu* operon expression *(i.e.,* 1,2-PD not added to growth media) only exhibit diffuse GFP (Figure S3). Both the ΔA and ΔJ single knockout strains appear capable of forming MCPs as indicated by the presence of puncta throughout the cytoplasm (Figure 3C). In the ΔA ΔJ strain, we primarily observed polar bodies indicative of protein aggregation rather than MCP formation (Figure 3C). Indeed, the number of fluorescent puncta in the double knockout strain was on average 1 punctum per cell – significantly lower than the average of 3-6 puncta per cell observed in the WT strain or the single knockouts (p < 0.0001) (Figure 3D). The negative control ΔpocR strain, in which the positive regulatory protein pocR for the *pdu* operon is knocked out so MCP proteins are not expressed, displayed mostly diffuse GFP fluorescence with occasional puncta (Figure 3C). The rare occurrence of puncta in this strain is consistent with published data indicating that signal sequences decrease protein solubility and occasionally lead to aggregation [44]. Thus, the occasional puncta in this strain are likely aggregated GFP. The ΔA ΔJ strain also had significantly more puncta than the ΔpocR negative control (p < 0.0001) but significantly fewer than the WT or single knockouts (Figure 3C, 3D). Cumulatively, these results imply that when MCPs do not assemble properly, induction of the *pdu* operon leads to more aggregation than induction of the tagged GFP constructs alone. This is likely due to the expression and aggregation of other *pdu* proteins, including enzymes and shell proteins, into a “proto-MCP” core [33,45]. These loose aggregates are then either engulfed by self-assembling shell proteins and form intact MCPs (as is the case for the WT, ΔA, and ΔJ strains), or they continue to aggregate and form polar bodies (as is the case for the ΔA ΔJ strain).

We also investigated the impact of shell modification on the function of MCP to metabolize 1,2-PD. The ΔA ΔJ strain had a variable growth pattern indicative of malformed MCPs when grown in media with 1,2-PD as the sole carbon source and supplemented with high (150 nM) adenosylcobalamin (AdoB_12_) (Figure S4A). The AdoB_12_ supplementation is necessary as the cofactor is required in the first step of 1,2-PD metabolism, but *S. enterica* cannot produce AdoB_12_ during aerobic growth. At high AdoB_12_ concentrations, malformed shells initially allow rapid consumption of 1,2-PD since the encapsulated enzymes are more accessible by the substrate, leading to rapid growth [23,24,38]. At the same time, however, the toxic intermediate propionaldehyde quickly accumulates, leaks out into the cytoplasm, and causes DNA damage that leads to stalled growth [23–25,38]. The ΔA ΔJ double knockout strain outgrew the WT and ΔA strains initially, but then growth stalled for several hours before continuing, indicating an initial build-up and subsequent loss of propionaldehyde (Figure S4A). This pattern of accumulation and loss of propionaldehyde was also seen in MCPs with PduA point mutations [24,25]. Stalled growth in the ΔJ strain is consistent with previous reports [38,42], but surprisingly, the stalled growth for the ΔA ΔJ strain was shorter than the ΔJ strain. It is important to note that, in addition to shell assembly defects, differences in shell protein function could also cause a growth defect. Despite their structural similarity, PduA and PduJ are thought to have functional differences with regard to metabolite diffusion [20]. Thus, while impaired MCP formation is a plausible cause for the growth defect for the ΔA ΔJ double knockout strain, the functional difference of PduA and PduJ could play a role in the growth difference between the ΔA and ΔJ single knockout strains. Subtle changes to enzyme encapsulation levels, substrate or cofactor diffusion, or MCP size could also play a role in cell growth and should be the subject of future investigations.

The results of these *in vivo* and *in vitro* assays suggest that knocking out *pduA* and *pduJ* together impairs MCP formation and function. While the outcomes of SDS-PAGE, TEM, and the GFP encapsulation assay all indicate that the ΔA ΔJ strain is unable to form MCPs, the results of the growth assay indicate subtle functional differences among the various knockout strains.

### Substitution of PduA with other Pdu shell proteins does not abolish MCP assembly

As the single knockouts of *pduA* or *pduJ* did not prevent MCP formation, we next investigated whether increased expression of other Pdu shell proteins could impact MCP formation. We chose to insert the other shell proteins in the *pduA* locus as it is at the beginning of the operon, and gene expression level is higher at the start than toward the end of the operon [46]. Furthermore, the use of the same position controls for any effect that genetic positioning within the operon may have on the function of the shell protein, as suggested by Chowdhury *et al.* [20]. To assess the effect of shell protein substitution, we replaced the *pduA* gene in the *pdu* operon with *pduK, -N, -U, -T,* or *-J* gene (ΔA::X) using the method of recombineering by Court (Figure 4A, Table 1) [47]. We also created a strain with *pduA* at the *pduJ* locus (ΔJ::A) to evaluate the effect of increasing the amount of *pduA* coded on the operon (Figure 4A, Table 1). The ability to manipulate shell expression and composition will be a valuable tool in the future efforts to reengineer MCPs for metabolic engineering.

**Figure 4.**
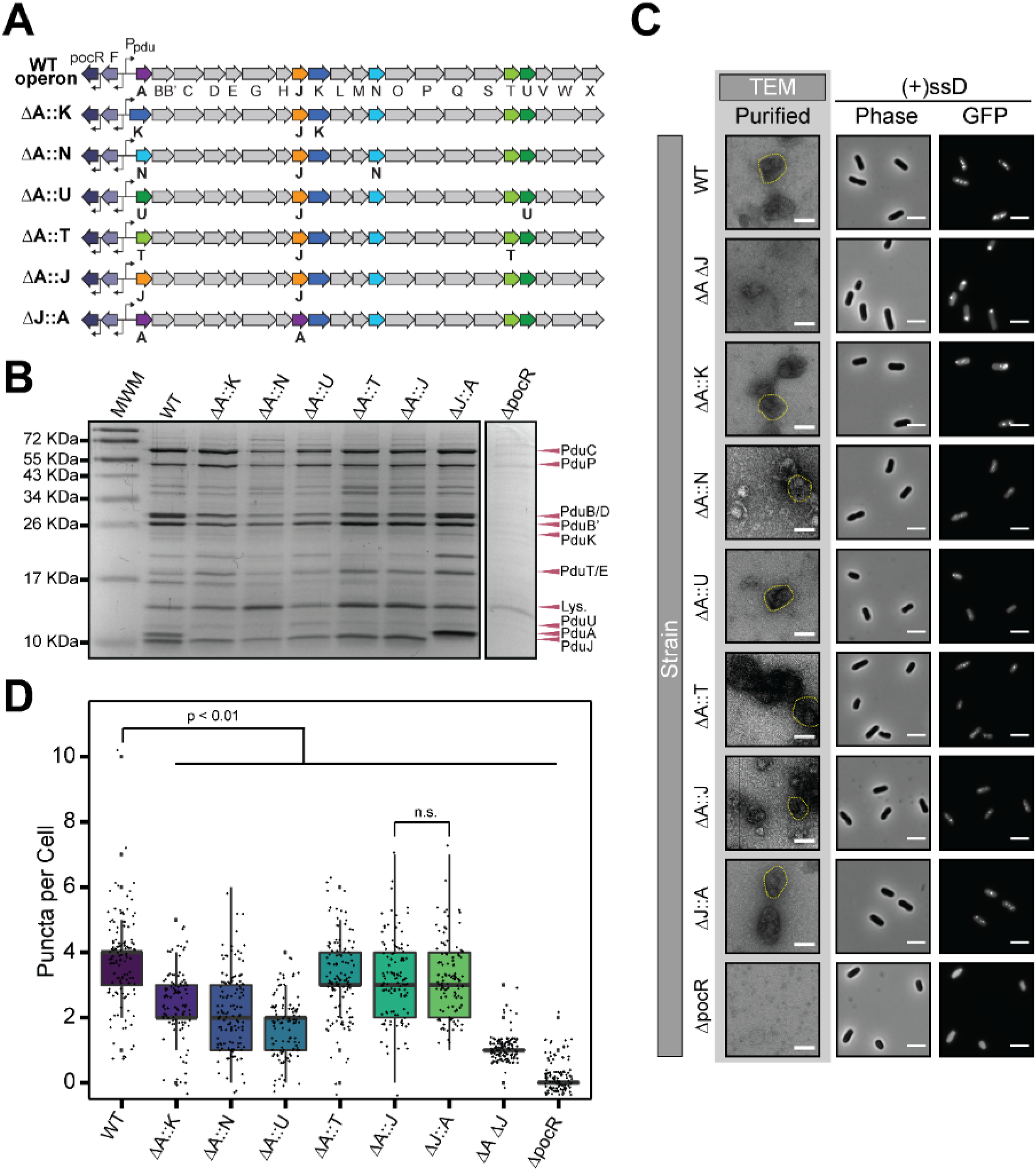
MCPs assemble in single knockout of pduA or pduJ substituted with other Pdu shell proteins. (A) Schematic representation of *pdu* operon, and *pdu* gene substitutions in *pduA* or *pduJ* single knockout strains. (B) Coomassie-stained SDS-PAGE gel of purified MCPs from single knockout substitution strains with Pdu MCP components labelled. “MWM” is the molecular weight marker and “Lys.” is lysozyme from the lysis buffer. (C) (Left) TEM of purified MCPs from single knockout substitution strains. Example MCPs are outlined in yellow. Scale bar = 100 nm. (Right) Phase contrast and GFP fluorescence microscopy images of strains encapsulating ssD-GFP. Row labels indicate the strain and column labels indicate the type of microscopy. Scale bars = 3 μm. (D) Quantification of puncta per cell for strains encapsulating ssD-GFP. WT had significantly higher number of puncta than the substitution strains, ΔA ΔJ strain, or ΔpocR strain (p < 0.01, pairwise t-test with Holm’s adjustment), but ΔA::J and ΔJ::A strains showed no significant difference between them (p = 1, pairwise t-test with Holm’s adjustment). Counts were collected from three biological replicates (>24 cells counted per strain per replicate). Overlaid data points represent the number of puncta for an individual cell and are distributed randomly around the whole number value (y-axis) to facilitate visualization.

**Table 1.** Strains used in this study.

| **Strain** | **Organism** | **Genotype** |
| --- | --- | --- |
| DTE017 | *E. coli* DH10b | Wild type |
| DTE001 | *E. coli* BL21 | Wild type |
| MFSS044 | *S. enterica* serovar Typhimurium LT2 | Wild type |
| TUC01 [47,66] | *E. coli* W3110 | *gal490 pglΔ8 λcl857 Δ(cro-bioA) int<->cat/sacB* |
| MFSS375 | *S. enterica* serovar Typhimurium LT2 | *ΔpduA* |
| NWKS081 | *S. enterica* serovar Typhimurium LT2 | *ΔpduJ* |
| NWKS083 | *S. enterica* serovar Typhimurium LT2 | *ΔpduA ΔpduJ* |
| SPIS051 | *S. enterica* serovar Typhimurium LT2 | *ΔpocR* |
| SPIS188 | *S. enterica* serovar Typhimurium LT2 | *ΔpduA::pduK* |
| MFSS409 | *S. enterica* serovar Typhimurium LT2 | *ΔpduA::pduN* |
| SPIS132 | *S. enterica* serovar Typhimurium LT2 | *ΔpduA::pduU* |
| MFSS411 | *S. enterica* serovar Typhimurium LT2 | *ΔpduA::pduT* |
| MFSS412 | *S. enterica* serovar Typhimurium LT2 | *ΔpduA::pduJ* |
| SPIS201 | *S. enterica* serovar Typhimurium LT2 | *ΔpduA::EutM* |
| MFSS387 | *S. enterica* serovar Typhimurium LT2 | *ΔpduA::csoS1A* |
| SPIS194 | *S. enterica* serovar Typhimurium LT2 | *ΔpduJ::pduA* |
| NWKS293 | *S. enterica* serovar Typhimurium LT2 | *ΔpduA::pduK ΔpduJ* |
| SPIS144 | *S. enterica* serovar Typhimurium LT2 | *ΔpduA::pduN ΔpduJ* |
| SPIS145 | *S. enterica* serovar Typhimurium LT2 | *ΔpduA::pduU ΔpduJ* |
| SPIS146 | *S. enterica* serovar Typhimurium LT2 | *ΔpduA::pduT ΔpduJ* |
| SPIS147 | *S. enterica* serovar Typhimurium LT2 | *ΔpduA::pduJ ΔpduJ* |
| SPIS189 | *S. enterica* serovar Typhimurium LT2 | *ΔpduA ΔpduJ::pduA* |
| NWKS262 | *S. enterica* serovar Typhimurium LT2 | *ΔpduA::pduA-K26A ΔpduJ* |
| NWKS294 | *S. enterica* serovar Typhimurium LT2 | *ΔpduA::pduJ-K25A ΔpduJ* |
| NWKS253 | *S. enterica* serovar Typhimurium LT2 | *ΔpduA::eutM ΔpduJ* |
| NWKS254 | *S. enterica* serovar Typhimurium LT2 | *ΔpduA::csoS1A ΔpduJ* |

First, we tested whether MCPs could be purified from the modified strains. We successfully purified MCPs from each of the variants and their banding patterns on the Coomassie-stained SDS-PAGE gel were similar to WT MCPs except for the expected missing band for the substituted PduA or PduJ at ~10 kDa (Figure 4B). Interestingly, we did not see an obvious increase in the intensity of the band corresponding to the shell protein that was duplicated at the *pduA* locus (Figure 4B). It should be noted that due to the limited number of vertices possessed by an MCP shell, the concentration of the pentameric cap PduN is typically too low to visualize on an SDS-PAGE gel [38]. TEM of the purified variant MCPs showed polyhedral structures with angular edges, as we typically observe with a WT Pdu MCP (Figure 4C).

To determine whether the shell gene substitutions impact the native pathway performance, we evaluated their growth in media with 1,2-PD and 150 nM AdoB_12_, as we did for the single and double knockouts of *pduA* and *pduJ.* All the ΔA::X strains except for the ΔA::U strain grew similarly (Figure S4B). The ΔA::U strain had a longer initial lag than the other strains and the greatest increase in doubling time (td) compared to the WT (Table S1) (p < 0.0001). We suspect the lag in the growth might be due to a transcription or translation issue, leading to delayed MCP formation, as we also observed a longer initial lag when we grew the strain with a limiting (20 nM) level of AdoB_12_ (Figure S4B insert). In media with limiting AdoB_12_, malformed MCPs can continue their rapid consumption of 1,2-PD without accumulating propionaldehyde to toxic levels, enabling these strains to outgrow the WT strain [24]. Instead, ΔA::U had lower cell density than WT for a large portion of the growth in the limiting AdoB_12_ condition (Figure S4B insert). The lack of dependence on AdoB_12_ level suggests that the ΔA::U growth defect is primarily due to reasons other than malformed MCPs.

We also evaluated the impact of Pdu shell substitution on MCP formation using the encapsulation assay with ssD-GFP (Figure 4C). As expected, we observed puncta when 1,2-PD was added to induce compartment formation (Figure 4C) and only diffuse GFP when 1,2-PD was not added (Figure S5A). The ΔA::X and ΔJ::A modified strains all displayed a higher puncta count per cell than the ΔA ΔJ strain or the negative control ΔpocR strain (p < 0.0001) (Figure 4D), but a lower puncta count per cell compared to the WT strain (p < 0.003). The reduced puncta count in the shell substitution strains could indicate a polar effect of the gene substitution that was not apparent on the SDS-PAGE gel or the growth curves, with the exception of the ΔA::U strain. The ΔA::U strain exhibited the greatest reduction in the puncta per cell count and this reduction could be the cause of the growth defect observed in the growth assay (Figure S4B). With translation or transcription issues delaying MCP formation, we hypothesized that there would be fewer MCPs available to metabolize 1,2-PD, thereby decreasing carbon flux and overall cell growth. Overall, the results indicate that substituting *pduA* with other Pdu shell proteins still allows MCP formation but positioning other Pdu shell genes at the beginning of the operon may have some polar effects that alter growth or encapsulation, which can be further investigated in future studies.

### PduA and PduJ are not fully interchangeable and are not replaceable by other Pdu shell proteins

After confirming that MCPs could still form with other Pdu shell proteins substituted at the *pduA* locus, we next investigated whether these shell proteins could substitute for PduA and rescue MCP formation in the ΔA ΔJ double knockout strain. Given that genes closer to the start of the *pdu* operon are typically expressed to a higher degree, we hypothesized that placing these genes at the *pduA* locus could enable them to compensate for the loss of PduA and PduJ via increased expression [46]. This would also control for the potential effect of gene locus on shell protein function, which has been reported to have an effect at the *pduA* locus [20]. To this end, we knocked out the *pduJ* gene in each of the ΔA::X strains (ΔA::X ΔJ) to create the double knockout substitution strains (Figure 5A). We also created a ΔA ΔJ::A strain to determine whether PduA and PduJ are fully interchangeable with one another (Figure 5A, Table 1).

**Figure 5.**
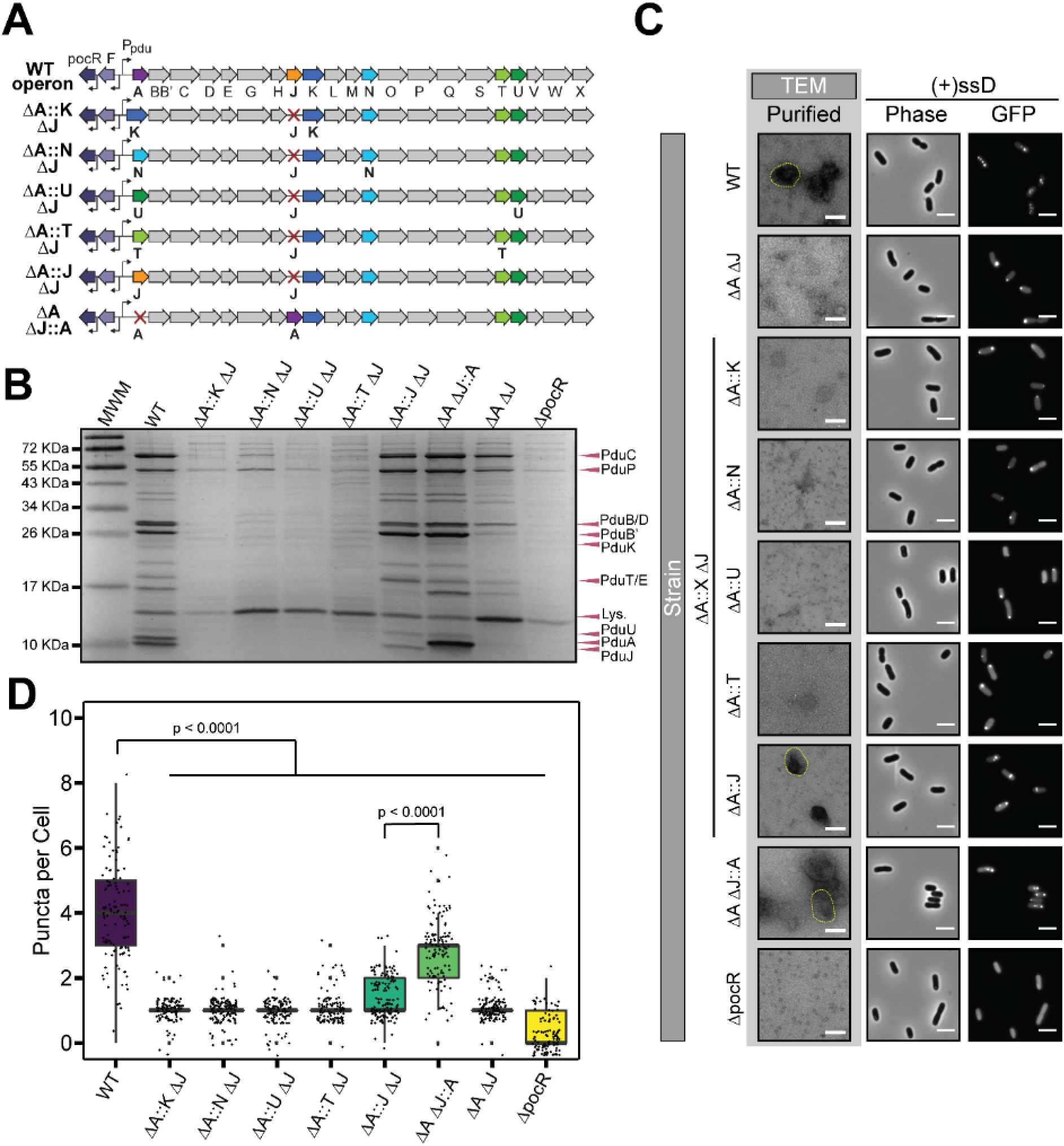
Only strains with pduA or pduJ form MCPs in double knockout substitution strains. (A) Schematic representation of *pdu* operon, and *pdu* gene substitutions in ΔA ΔJ double knockout strain. (B) Coomassie-stained SDS-PAGE gel of purified MCPs from double knockout substitution strains with Pdu MCP components labelled. “MWM” is the molecular weight marker and “Lys.” is lysozyme from the lysis buffer. (B) (Left) TEM of purified MCPs from double knockout substitution strains. Example MCPs are outlined in yellow. Scale bar = 100 nm. (Right) Phase contrast and GFP fluorescence microscopy images of strains encapsulating ssD-GFP. Row labels indicate the strain and column labels indicate the type of microscopy. Scale bars = 3 μm. (D) Quantification of puncta per cell for strains encapsulating ssD-GFP. WT had significantly higher number of puncta than substitution strains (p < 0.0001, pairwise t-test with Holm’s adjustment). ΔA ΔJ::A strain also had higher puncta count than the ΔA::J ΔJ strain (p<0.0001, t-test). Counts were collected from three biological replicates (>28 cells counted per strain per replicate). Overlaid data points represent the number of puncta for an individual cell and are distributed randomly around the whole number value (y-axis) to facilitate visualization.

Substitutions of other Pdu shell proteins at the *pduA* locus could not rescue MCP formation in the absence of both PduA and PduJ. We purified and imaged MCPs and conducted the encapsulation assay on the double knockout substitutions, as we did for the single knockout substitutions. Only MCPs purified from the ΔA::J ΔJ and ΔA ΔJ::A strains showed the banding pattern similar to WT MCPs on the SDS-PAGE gel (Figure 5B). The SDS-PAGE banding pattern for MCPs purified from the rest of the strains resembled the pattern for the ΔA ΔJ or the ΔpocR knockout strain. Similarly, TEM showed the typical MCP morphology only for samples from the ΔA::J ΔJ and ΔA ΔJ::A strains, and aggregates for samples from the rest of the modified strains (Figure 5C). To further confirm that other shell proteins cannot rescue MCP formation, we also overexpressed each shell protein (PduBB’, -K, -N, -T, and -U) from a plasmid in the ΔA ΔJ strain. We were unable to purify MCPs from these strains, as indicated by SDS-PAGE gel (Figure S6A). We conclude that other Pdu shell proteins are unable to rescue MCP formation even after overexpression, supporting our results from chromosomal modifications.

The results of the encapsulation assay bolster the analysis of the purified MCPs. Only the ΔA::J ΔJ and ΔA ΔJ::A variants gave rise to puncta counts higher than the ΔA ΔJ or ΔpocR strains in the samples induced for MCP formation (p < 0.0001) (Figure 5D). All strains displayed only diffuse GFP when they were not induced for MCP formation (Figure S5B). We counted more puncta per cell in the ΔA ΔJ::A variant than the ΔA::J ΔJ strain (Figure 5D). This was in contrast to the single substitution variants, in which the ΔA::J and ΔJ::A strains showed no significant difference in the puncta per cell count (Figure 4D). Taken together, these results confirm the prior report that PduA and PduJ are structurally redundant but are not fully interchangeable [20]. To our knowledge this is the first demonstration that at least one of these proteins is required for MCP formation and other Pdu shell proteins cannot substitute for PduA or PduJ to form MCPs in the absence of both PduA and PduJ.

### Assembly-deficient pduA and pduJ point mutants do not rescue compartment assembly

Once we established that the presence of PduA or PduJ is essential for MCP formation and cannot be replaced by other Pdu shell proteins, we then tested the hypothesis that self-assembly into tubes is a critical property for PduA and PduJ function and is linked to MCP formation. A conserved lysine residue at position 26 and 25 of PduA and PduJ, respectively, is critical for self-assembly into higher-order structures composed of multiple hexamers [42]. These conserved lysine residues are at the outer edge of the hexamers and form hydrogen bonds with adjacent hexamers in self-assembled structures [28]. Mutating these conserved lysine residues to alanine inhibits higher-order self-assembly while maintaining a folded, individual hexameric protein complex, as demonstrated in published crystallographic studies [20,42]. Taking advantage of this property, we integrated *pduA-K26A* and *pduJ-K25A* into the *pduA* locus in the *pduJ* deletion strain (Figure 6A). These strains contain only an assembly-deficient version of PduA or PduJ, allowing us to test the hypothesis that assembly-competent PduA or PduJ is essential for MCP formation.

**Figure 6.**
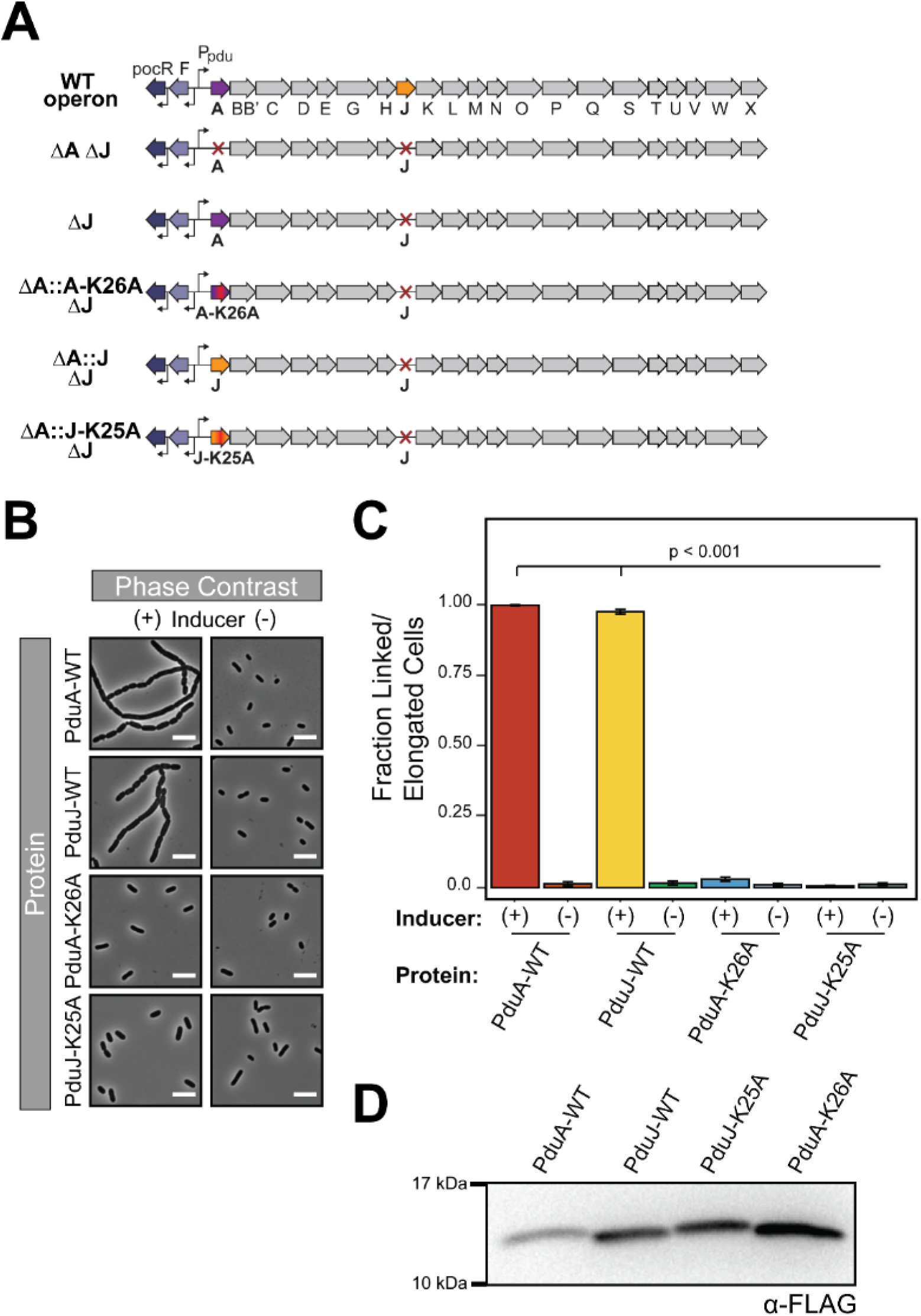
Point mutations disrupt PduA and PduJ self-assembly. (A) Schematic representation of *pdu* operon and strain modifications. (B) Phase contrast microscopy images of cells overexpressing PduA, PduJ, and assembly-deficient point mutants. (C) Fraction of linked or elongated cells in population of cells induced (+) or uninduced (-) for overexpression of WT or assembly deficient PduA and PduJ. Cells overexpressing (+ inducer) PduA-WT or PduJ-WT have a significantly greater fraction of cells in linkages of three or more cells or longer than 6 μm (p < 0.001, t-test). Counts were taken from three biological replicates (>80 cells counted per strain per replicate), and error bars represent standard error of the means. (D) Western blot of whole-cell lysate from cells overexpressing FLAG-tagged PduA, PduJ, and assembly-deficient point mutants.

We first verified that PduA-K26A and PduJ-K25A do not self-assemble into tubes by overexpressing these proteins in *E. coli* BL21 cells. Cells overexpressing these point mutants did not exhibit the cell division defect indicative of self-assembled proteins observed for the WT PduA and PduJ proteins (Figure 6B). While cells overexpressing WT PduA or PduJ were elongated or in chains of three or more cells over 90% of the time, less than 3% of cells overexpressing the assembly-deficient point mutants were elongated or in such chains (p < 0.001) (Figure 6C). This result did not appear to be due to a lack of expression of the assembly-deficient point mutants, as both mutants were detectable on a western blot of whole cell lysate from cells expressing the proteins (Figure 6D). Our results support previous findings by Pang and Frank *et al.,* which qualitatively demonstrated that mutating the conserved lysine disrupted PduA self-assembly into aberrant structures [39].

These well-characterized PduA-K26A and PduJ-K25A assembly-deficient mutants have only been evaluated in the native system when the other self-assembling protein (WT PduJ or PduA, respectively) is still present [42], potentially confounding results. Therefore, we tested the ability of these assembly-deficient mutants to rescue MCP formation in the absence of WT PduA or PduJ. Comparison of these strains allows us to test the hypothesis that only assembly-competent versions of PduA or PduJ enable MCP formation. MCPs were purified from strains with *pduA-K26A* and *pduJ-K25A* at the *pduA* locus and compared to MCPs purified from strains with WT *pduA* and *pduJ* at the same locus (Figure 6A). These four strains also had *pduJ* knocked out, since our results with ΔA::J ΔJ indicate that PduJ can compensate for the loss of a functional protein expressed from the *pduA* locus (Figure 5). On SDS-PAGE gels, only strains containing WT *pduA* (ΔJ) or *pduJ* (ΔA::J ΔJ) at the *pduA* locus displayed the expected banding pattern of assembled MCPs (Figure 7A). By contrast, strains containing either *pduA-K26A* or *pduJ-K25A* displayed a banding pattern similar to the ΔA ΔJ strain or the ΔpocR knockout strain (Figure 7A), indicative of protein aggregation. TEM confirmed that strains containing WT *pduA* or *pduJ* produced MCPs with WT morphology, while strains containing the assembly-deficient point mutants yielded mostly protein aggregates (Figure 7B). We also tested the possibility that overexpressing the mutant shell proteins could rescue MCP assembly. To this end, we overexpressed each mutant shell protein (PduA-K26A and PduJ-K25A) from a plasmid in the ΔA ΔJ strain. However, even after overexpression from a plasmid, the assembly-deficient mutants were unable to rescue MCP formation, and we were unable to purify MCPs from these strains, as indicated by SDS-PAGE gel (Figure S6B).

**Figure 7.**
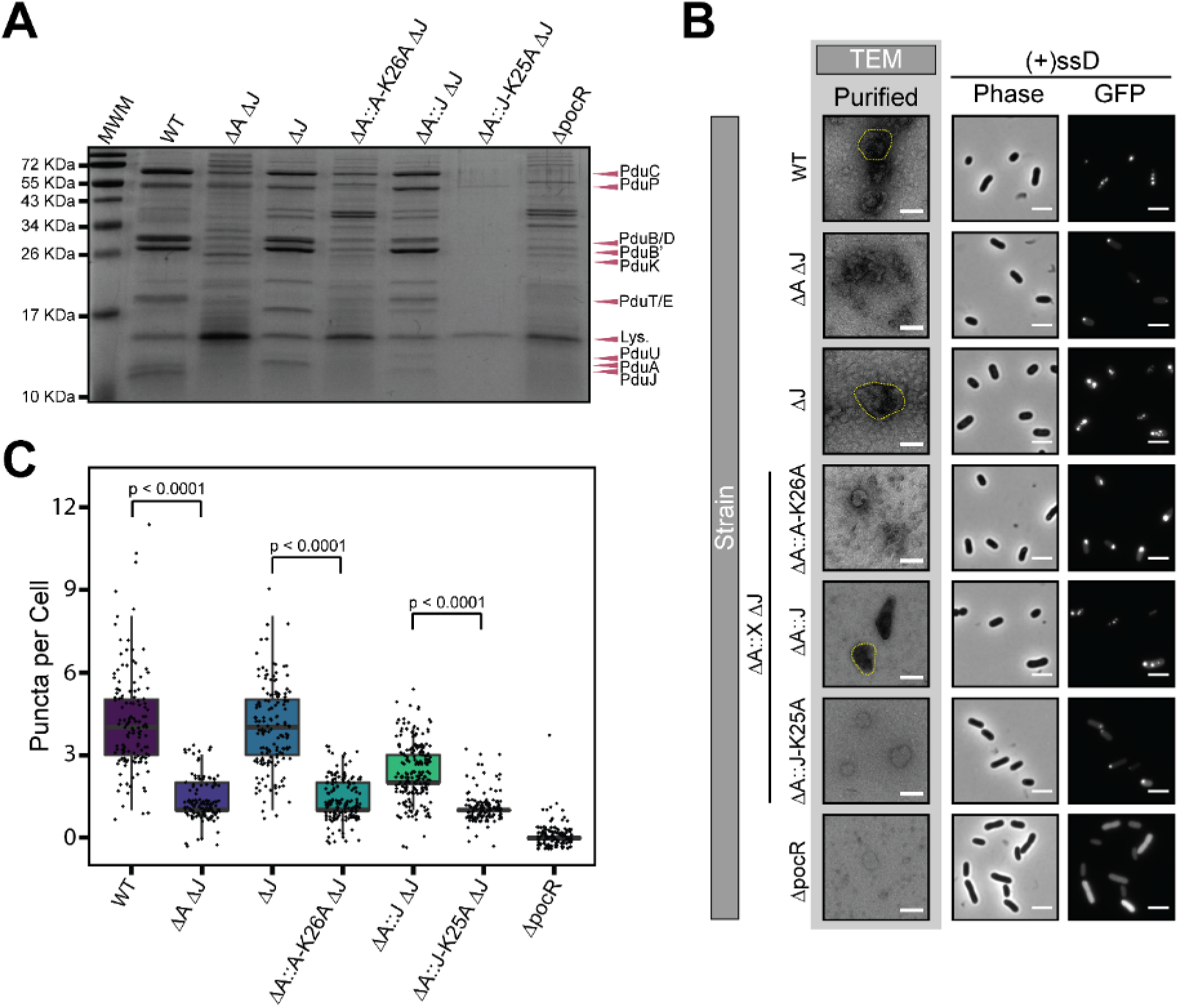
Assembly-deficient point mutants do not rescue MCP formation. (A) Coomassie-stained SDS-PAGE gel of purified MCPs from different strains. “MWM” is the molecular weight marker, and “Lys.” is lysozyme. (B) (Left) TEM of purified MCPs from different strains. Example MCPs are outlined in yellow. Scale bars = 100 nm. (Right) Phase contrast and GFP fluorescence microscopy images of strains encapsulating ssD-GFP. Row labels indicate the strain and column labels indicate the type of microscopy. Scale bars = 3 μm. (C) Quantification of puncta per cell for strains encapsulating ssD-GFP. Strains expressing assembly-deficient shell proteins had significantly fewer puncta per cell than strains expressing either PduA-WT or PduJ-WT (p < 0.0001, t-test). Counts were collected from three biological replicates (>40 cells counted per strain per replicate). Overlaid data points represent the number of puncta for an individual cell and are distributed randomly around the whole number value (y-axis) to facilitate visualization.

We further characterized MCP formation in these strains using the GFP encapsulation assay with ssD-GFP as described above. As expected, strains with WT *pduA* or *pduJ* at the *pduA* locus contained numerous bright fluorescent puncta throughout their cytoplasm when MCP expression was induced, indicative of proper MCP formation and GFP encapsulation (Figure 7B). Conversely, strains containing the assembly-deficient point mutants at the *pduA* locus displayed mostly polar bodies, similar to the ΔA ΔJ double knockout strain and indicative of protein aggregation rather than MCP formation (Figure 7B). Strains expressing PduA-K26A or PduJ-K25A showed significantly fewer puncta per cell (p < 0.0001) compared to strains expressing WT PduA or PduJ, implying that the loss of self-assembly indeed inhibits the function of these proteins in MCP assembly (Figure 7C). As with other strains in this work, only diffuse ssD-GFP fluorescence was observed when MCP expression was not induced (Figure S5C). These results support our hypothesis that self-assembly is an important property for the native function of PduA and PduJ and is associated with overall MCP assembly, building on reports by Sinha *et al.* and Pang and Frank *et al.* [39,42].

### PduA and PduJ homologs do not rescue microcompartment assembly

Self-assembly upon overexpression is a unique property of PduA and PduJ within the *pdu* operon. However, PduA homologs from other microcompartment systems can also self-assemble. For example, EutM from the ethanolamine utilization (Eut) microcompartment self-assembles into long tubes, similar to PduA [48,49]. Therefore, we hypothesized that PduA homologs might be able to rescue MCP formation, so we integrated the PduA homologs *eutM* and *csoS1A* (from a carboxysome operon in *Halothiobacillus neapolitanus)* into the *pduA* locus. These two proteins contain BMC-domains and are structurally similar to PduA and PduJ, with small differences (Figure 8A).

**Figure 8.**
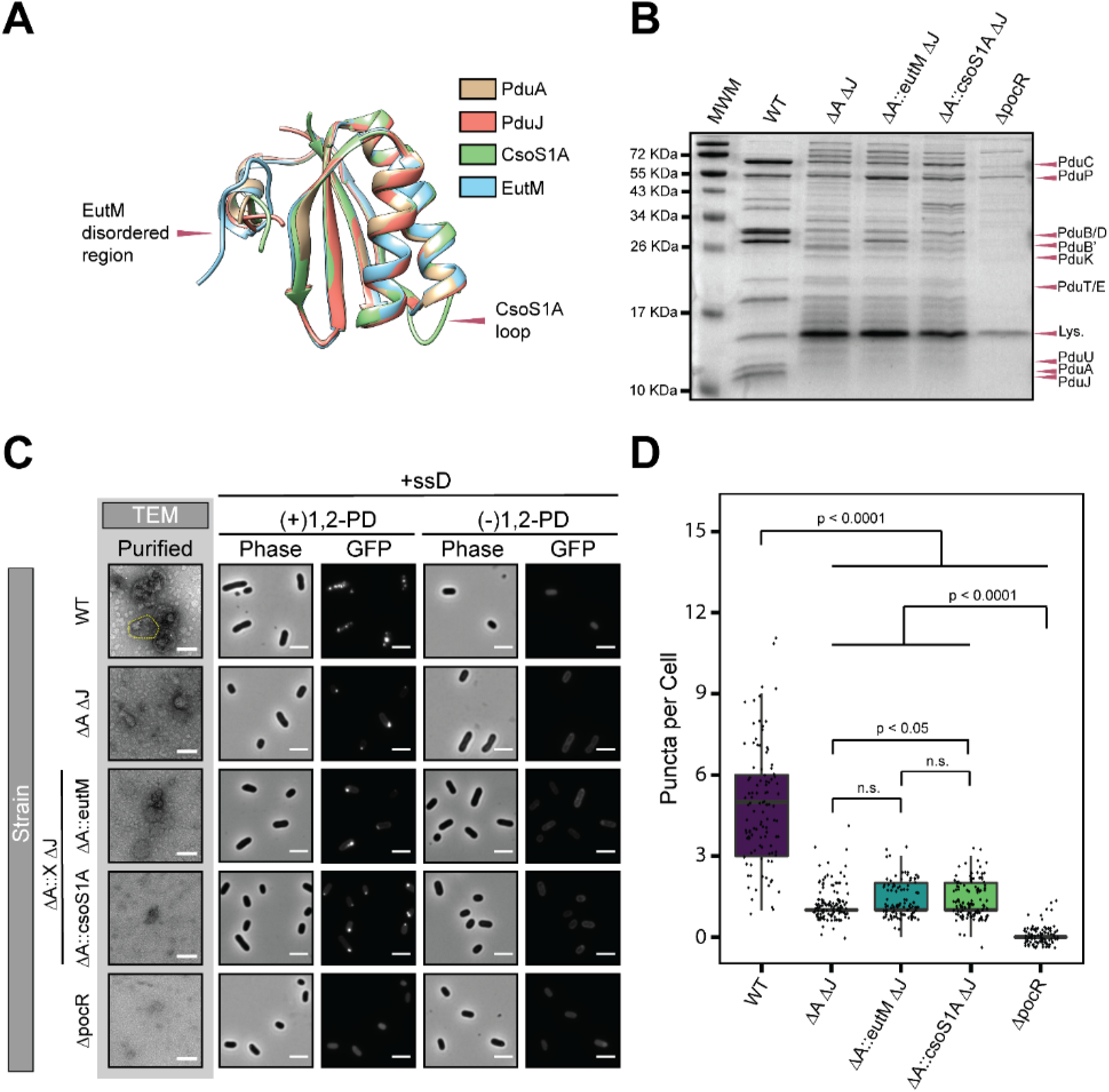
PduA homologs do not rescue MCP formation. (A) Structural overlay of PduA (tan, PDB ID: **<u>3NGK</u>**) [19], PduJ (pink, PDB ID: **<u>5D6V</u>**) [20], EutM (blue, PDB ID: **<u>3MPY</u>**) [63], and CsoS1A (green, PDB ID: **<u>2EWH</u>**) [64]. Structures were visualized using Chimera [65]. (B) Coomassie-stained SDS-PAGE gel of purified MCPs. Strains are listed above lanes on the gel. “MWM” is the molecular weight marker and “Lys.” is lysozyme. (C) (Left) TEM of purified MCPs from different strains. Scale bars = 100 nm. (Right) Phase contrast and GFP fluorescence microscopy images of strains encapsulating ssD-GFP. Row labels indicate the strain and column labels indicate the type of microscopy. Scale bars = 3 μm. (D) Quantification of puncta per cell for strains encapsulating ssD-GFP. Strains expressing EutM or CsoS1A have significantly fewer puncta per cell than the wild-type (WT) strain (p < 0.0001, t-test). Counts were collected from three biological replicates (>33 cells counted per strain per replicate). Overlaid data points represent the number of puncta for an individual cell and are distributed randomly around the whole number value (y-axis) to facilitate visualization.

The SDS-PAGE banding pattern of MCPs purified from strains containing *eutM* or *csoS1A* at the *pduA* locus and lacking *pduJ* appeared similar to that of the ΔA ΔJ strain, indicative of poorly-formed MCPs or protein aggregation (Figure 8B). Indeed, TEM of purified MCPs from these strains revealed primarily aggregated proteins (Figure 8C). Further, overexpressing these proteins from a plasmid also did not rescue MCP formation, as indicated by the SDS-PAGE banding pattern (Figure S6C). The GFP encapsulation assay further confirmed these results. The ΔA::eutM ΔJ and ΔA::csoS1A ΔJ strains contained mostly polar bodies (Figure 8C), similar to the ΔA ΔJ double knockout strain, and the substituted strains had significantly fewer puncta per cell than the WT strain (p < 0.0001) (Figure 8D). Strains in which MCPs were not induced displayed only diffuse GFP fluorescence (Figure 8C). Combined, these results imply that Pdu MCP formation relies specifically on PduA or PduJ, as substitution of other BMC-domain proteins was unable to produce intact MCPs. Interestingly, we showed that EutM and CsoS1A do not seem to self-assemble to the same extent as PduA and PduJ, as determined by the cell elongation assay, in spite of similar expression levels (Figure S7). We suspect that EutM and CsoS1A are capable of self-assembly, but not to the extent that they form long tubes detectable in our microscopy-based assay. Moreover, we hypothesize that they are unable to interact properly with other members of the Pdu MCP shell or enzymatic core, impairing their ability to rescue MCP function. This would support findings that EutM can be incorporated into the Pdu MCP shell, but is perhaps different enough that it cannot drive Pdu MCP assembly on its own, without the presence of PduA or PduJ [25]. This idea of incorporation, but not complete functional complementation, is also supported by studies done in carboxysomes, where an α-carboxysome shell protein was incorporated but unable to rescue β-carboxysome function [50].

## Discussion

MCPs are intriguing prokaryotic organelles that are implicated in the infectivity of certain human pathogens [51–57]. MCPs also offer a unique opportunity to address shortcomings in metabolic engineering, while providing a model system to study protein self-assembly. This incredible breadth of scope, ranging from basic understanding of protein science to infectious diseases, necessitates the development of proper controls. In this work, we uncovered an essential part of the MCP shell assembly mechanism and in the process also developed a new control for Pdu MCP shell assembly. Our findings have the potential to impact how MCPs are studied in the future.

To thoroughly and quantitatively investigate the self-assembly of two Pdu MCP shell proteins, PduA and PduJ, we developed a novel light microscopy-based method to rapidly quantify the self-assembly property without protein purification. While runaway self-assembly of PduA into tubes has been demonstrated previously [39], our work presents a rapid, alternative to EM-based methods. Lee *et al.* used tomography on an engineered PduA homolog from *Citrobacter freundii* to assess self-assembly, but this method requires extensive sample preparation and computational analysis, and likely underestimates the extraordinary length of these structures due to the size limitations of ultra-thin sections [40]. Our light microscopy-based method enables quantification of changes to PduA or PduJ self-assembly into extended tubes without the need for purification of the protein assemblies or modifying the native *pdu* operon. Visualization and quantification of self-assembly is rapid and straightforward, as cell division defects are easily observable under a light microscope and do not require specialized or complex software for analysis. This technique also has potential for utility outside of the Pdu MCP system in assessing assembly of other fibril-forming proteins without the need for protein purification.

The self-assembly property of PduA and PduJ appears to be a conserved trait among BMC-domain proteins from other systems. EutM in the Eut microcompartment system and RmmH from a mycobacterium operon both assemble into tubes upon overexpression [41,48,49]. Heterologous expression of carboxysome shell proteins also leads to fibril formation at high concentrations [58]. Thus, we hypothesized that a high propensity towards self-assembly is an essential feature for MCP assembly. Indeed, we showed that PduA and PduJ are essential and redundant for shell assembly, and that this is linked to their ability to individually self-assemble. It is possible that an “assembler” (*i.e.,* a protein that is prone to assembly into large structures at high concentrations) is necessary for all MCP systems. The existence of an “assembler” protein would allow other BMC domain-containing shell proteins to have specific functions apart from shell formation. For example, in the Pdu MCP system, PduB is important for enzyme encapsulation [32] while PduT may play a role in electron transfer across the shell [19,29]. These proteins might only need to retain an ability to interact with the “assembler” to be incorporated into the shell, instead of requiring a self-assembling nature of their own (Figure 9). This fits together well with current models of MCP shell assembly, in which the shell either assembles before or concomitantly with the enzymatic core (Figure 9) [59]. We propose that spontaneous shell assembly is initiated by the “assembler” proteins (such as PduA and PduJ in the Pdu system), and they help connect the other shell proteins and the core (Figure 9). This interaction with other shell proteins, specifically PduN, likely helps impart the MCP shape and promotes assembly into flat structures, rather than the curved tubes. This is supported by *pduN* knockouts, which formed elongated structures rather than the normal, polyhedral MCPs [38]. Assembly with PduB also likely flattens the shape of the assembled structures, as the interaction between trimers and hexamers occurs at a lower angle (25° vs. 30°) [50], and PduB itself assembles into structures with wider diameters [35]. Interaction with the core, either directly through PduA and PduJ [26,27], or indirectly through PduB [32] or PduK [12], also likely plays a role in imparting the final polyhedral structure. Identification of the essentiality of a self-assembling protein will facilitate future investigations into the function of other MCP shell proteins as well as for the creation of a new minimal, engineered Pdu MCP with fewer shell proteins than currently published in literature [36].

**Figure 9.**
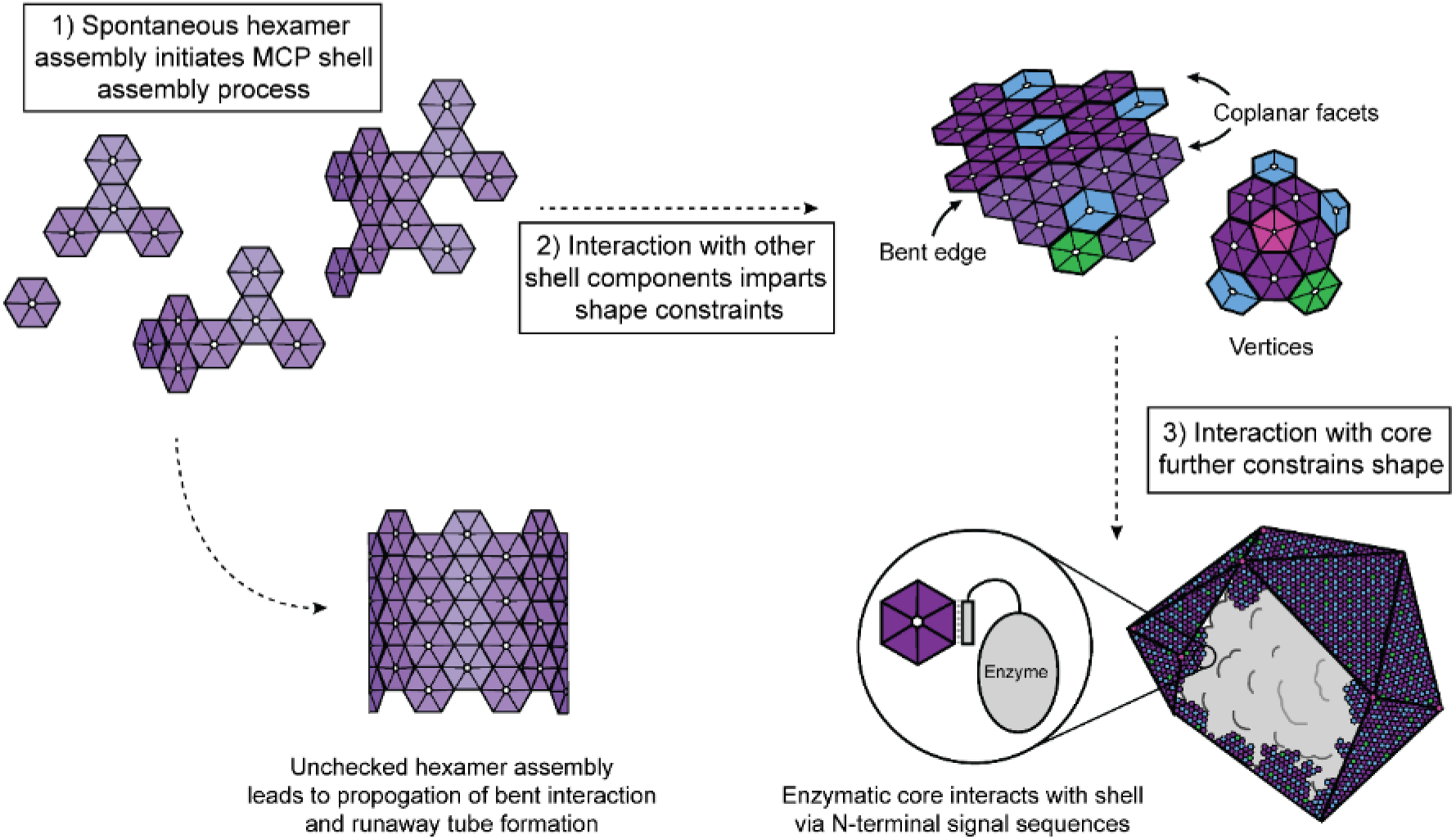
PduA and PduJ assembly is necessary for the Pdu MCP assembly mechanism. Schematic representation of Pdu MCP assembly, in which PduA and PduJ spontaneously assemble to initiate MCP assembly. PduA and PduJ mediate interactions with other shell proteins and the enzymatic core. These additional interactions promote assembly into the final MCP architecture.

In the process of uncovering the roles of PduA and PduJ self-assembly in MCP formation, we discovered that the ΔA ΔJ strain forms protein aggregates but not intact MCPs. The inability of other Pdu shell proteins to compensate for the ΔA ΔJ double knockout demonstrated the importance of having a shell protein with the self-assembling property. The only strains that rescued MCP formation of the ΔA ΔJ double knockout strain were those with *pduJ* substituted at the *pduA* locus or *pduA* substituted at the *pduJ* locus. None of the other tested Pdu shell proteins, despite having a BMC-domain like PduA and PduJ, could form MCPs when substituted at the *pduA* locus. Thus, simply having a BMC-domain in the structure is insufficient to rescue MCP formation. Furthermore, duplicating and moving a copy of a shell protein gene upstream to the start of the *pdu* operon does not appear to affect MCP formation or shell protein incorporation significantly. For example, PduT and PduU shell proteins are thought to compose a smaller fraction of the Pdu shell than other shell proteins since they are encoded toward the end of the *pdu* operon [8]. However, the ΔA::X substitution of the tested shell proteins, including ΔA::T and ΔA::U, resulted in no obvious increase in the intensity of the corresponding band on the SDS-PAGE gel. A more quantitative (*e.g*., using mass spectrometry) evaluation of the ratio of MCP proteins in the single substitution variants would be a valuable future avenue of exploration [22]. Understanding how to control shell composition by manipulating the operon will be an important tool for engineering the Pdu MCP shell for metabolic engineering efforts, particularly if altered transport of metabolites is required.

The findings in this work, and in particular the ΔA ΔJ strain as a new non-assembly control, have broad applicability and utility in future investigations of the native Pdu MCP function as well as in the development of Pdu MCPs as nanobioreactors. For example, the ΔA ΔJ strain will facilitate studies that require a control with unlimited substrate diffusion across the shell. Inclusion of a shell-less control also will allow comparison of encapsulated and non-encapsulated pathways, enabling evaluation of the benefits of encapsulating heterologous pathways.

The necessity of the self-assembly property and the redundancy of PduA and PduJ revealed in this work will likely influence how Pdu and other similar MCPs are engineered. Consideration of the shell protein self-assembly will be a necessary part of the future design efforts. The impact of chromosomal position on the encapsulation assay we saw with the ΔA ΔJ::A variant versus the ΔA::J ΔJ strain also highlights the importance of being considerate of how and where these operons are manipulated. These results in our work, in addition to providing a useful control, provide a piece of the mechanistic puzzle for the assembly of these complex protein structures.

## Materials and Methods

### Plasmid and strain construction

The strains used in this study are listed in Table 1. All plasmids were constructed using Golden Gate cloning (Table S2) [60]. Briefly, inserts containing BsaI cut sites at the 5’ and 3’ ends were amplified using PCR with primers listed in Table S3. These inserts were cloned into an arabinose-inducible, Golden Gate-compatible vector with a p15a origin of replication and a chloramphenicol selectable marker (pBAD33t). FastDigest Eco31I from Thermo Fisher Scientific and T4 DNA ligase from New England Biolabs, Inc. were used for the Golden Gate reaction mixture. The ligations were transformed into *E. coli* DH10b cells and successful cloning was confirmed via Sanger sequencing. For overexpression tests, plasmids were transformed into *E. coli* BL21 cells for overexpression tests. Point mutants of PduA and PduJ were constructed using site-directed mutagenesis and KOD Hot Start polymerase (MilliporeSigma).

The *pdu* operon of *Salmonella enterica* serovar Typhimurium LT2 was modified using the λ Red recombineering method. Briefly, a *cat/sacB* selection cassette was first amplified using PCR to contain homologous 5’ and 3’ overhangs matching the target chromosomal location (Table S3). Using the recombineering method of the Court Lab [47], the selectable *cat/sacB* marker was inserted into the desired genomic location and then replaced with either a PCR-amplified gene of interest containing 5’ and 3’ homology to the target locus, or was knocked out, effectively removing the native gene. To reduce the potential polar effects from full deletion of a gene, the C-terminal 41 – 51 base pairs of *pduA* and *pduJ* loci were retained in each modification. The modified loci were sequence confirmed via PCR amplification and Sanger sequencing.

### Shell protein overexpression

Single colonies from cells streaked on agar plates were inoculated into 5 mL Lysogeny Broth, Miller (LB-M, Thermo Fisher Scientific) media in 24-well blocks. The blocks were sealed with Andwin Scientific AeraSeal™ film and grown overnight at 37 °C, 225 RPM for 15-16 hours. Overnight cultures were subcultured 1:100 (50 μL) into 5 mL of LB-M and grown at 37 °C, 225 RPM. Expression was induced with 0.2% (w/v, final concentration) L-(+)-arabinose 90 minutes after subculture, which typically corresponded to an OD_600_ of ~0.3. Cells were grown at 37 °C, 225 RPM for a minimum of 4 hours post-induction and then imaged under the microscope (see *Phase contrast and fluorescence microscopy).* The LB-M cultures and agar plates included 34 ug/mL chloramphenicol.

### GFP encapsulation assay

Overnight cultures of *S. enterica* LT2 strains containing pBAD33t-ssD-GFPmut2 plasmid (CMJ069), were prepared as described above into fresh 5 mL LB-M cultures in 24-well blocks and grown for 15-16 hours. The overnight cultures were then subcultured 1:500 (10 μL) into 5 mL of LB-M with 0.02% (w/v, final concentration) L-(+)-arabinose, 34 ug/mL chloramphenicol, and 0.4% (v/v, final concentration) 1,2-PD. Negative controls, where compartments were not induced, were grown without 0.4% (v/v) 1,2-PD in the growth media. Cultures were grown at 37 °C, 225 RPM for a minimum of 6 hours post-subculture and then imaged under the microscope (see *Phase contrast and fluorescence microscopy*).

### Growth assay

Single colonies of the WT and chromosomally modified *S. enterica* LT2 strains were inoculated into 5 mL of Terrific Broth media (Dot Scientific, Inc.) without glycerol in test tubes and grown for 15-16 hours at 37 °C, 225 RPM. The overnight cultures were subcultured to OD_600_ of 0.05 in 50 mL of Non-Carbon E (NCE) media (29 mM potassium phosphate monobasic, 34 mM potassium phosphate dibasic, 17 mM sodium ammonium hydrogen phosphate) supplemented with 1 mM magnesium sulfate, 50 μM ferric citrate, 55 mM 1,2-PD, and 150 nM or 20 nM adenosylcobalamin (Santa Cruz Biotechnology). The cultures were grown in 250 mL Erlenmeyer flasks at 37 °C, 225 RPM. OD_600_ measurements were taken on BioTek Synergy HTX multi-mode plate reader every 3 hours using 200 μL of sample in a clear, flat-bottom 96-well plate. From 12 hours and on, samples were diluted 1/5 or 1/10 in the culture media prior to measurement to remain within the linear range of the instrument. At least three biological replicates were measured for each strain. The doubling time (td) was calculated by td = log(2)/r, where r is the slope of linear region in the plot of Log(OD_600_) versus time.

### MCP expression and purification

For MCP expression, single colony inoculations of *S. enterica* LT2 strains were prepared in 5 mL LB-M and grown for 24 hours at 30 °C, 225 RPM. These saturated overnight cultures were used at 1:1000 ratio to inoculate large-scale expression cultures (200 mL NCE media supplemented with 1 mM magnesium sulfate, 50 μM ferric citrate, 42 mM succinate, and 55 mM 1,2-PD). Expression cultures were grown in 1 L Erlenmeyer flasks at 37 °C, 225 RPM until cultures reached an OD_600_ of at least 1.0 (usually 16-24 hours). In experiments where shell proteins were overexpressed, 34 ug/mL (final concentration) chloramphenicol was added, and cultures were induced at an OD_600_ = 0.2 with 0.02% (w/v, final concentration) L-(+)-arabinose.

MCPs were purified via differential centrifugation as previously reported [61]. Briefly, once expression cultures reached the target OD_600_, cells were pelleted at 5000 RPM, 4 °C, and lysed chemically using a Tris-buffered lysis solution (32 mM Tris, 200 mM potassium chloride, 5 mM magnesium chloride, 1.2% (v/v) 1,2-PD, 0.6% (w/v) octylthioglucoside (OTG), 2.2 mM β-mercaptoethanol, 0.8 mg/mL lysozyme (Thermo Fisher Scientific), and 0.04 U/mL DNase I (New England Biolabs, Inc.)). Cell lysate was clarified via centrifugation at 12,000 x *g,* 4 °C, for 5 minutes using a FIBER*Lite* F10BCI-6×500y rotor (Piramoon Technologies) in an Avanti JXN-30 centrifuge (Beckman Coulter). The clarified cell lysate was further centrifuged in a JS-24.38 rotor (Beckman Coulter) at 21,000 x *g*, 4 °C, for 20 minutes to pellet the microcompartments. The pelleted microcompartments were stored in a buffer of 50 mM Tris (pH 8.0), 50 mM potassium chloride, 5 mM magnesium chloride, 1% (v/v) 1,2-PD at 4 °C. The purified MCPs were fixed and prepared for TEM within 10 days of purification. Protein concentration was determined using the Pierce^TM^ BCA Protein Assay Kit (Thermo Fisher Scientific) with bovine serum albumin as standard.

### Phase contrast and fluorescence microscopy

Phase contrast and fluorescence microscopy was done on cells using Fisherbrand™ frosted microscope slides (Thermo Fisher Scientific Cat# 12-550-343) and 22 mm x 22 mm, #1.5 thickness cover slips (VWR Cat# 16004-302). Cells were imaged using a Nikon Eclipse Ni-U upright microscope and 100X oil immersion objective. GFP fluorescence was observed using the C-FL Endow GFP HYQ bandpass filter and collected with exposure time of 80 ms for strains with ssD-GFPmut2 strains and 100 ms for strains with ssP-GFPmut2. Digital micrographs were collected using an Andor Clara digital camera and NIS Elements Software (Nikon). All images were equally adjusted, within experiments, for brightness and contrast adjusted using ImageJ software [62]. Cell length measurements were collected using the segmented line tool in ImageJ [62]. Cell counts were collected on brightness and contrast-adjusted images. If a clear cleavage furrow was present on a contiguous cell body, the space flanking each furrow was counted as a single cell. For example, if a cell body contained three noticeable furrows (indicating an attempted cell-division event), this was counted as four linked cells. If a cell body was >10 μm in length, but no cleavage furrows were noticeable, the cell was counted as a single “linked” cell in the calculation for “fraction linked cells” (see Figure 1E, 6B).

### Ultra-thin section transmission electron microscopy

The bacterial cell samples were pelleted and fixed in a solution of 2.5% Electron Microscopy (EM) grade glutaraldehyde and 2% paraformaldehyde in 0.1 M 1,4-piperazinediethanesulfonic acid (PIPES) buffer. The samples were post fixed with 1% osmium tetroxide and dehydrated in a graded series of ethanol before infiltration and embedment with EMBed 812 epoxy resin. The epoxy resin was then cured at 60 °C for 48 hours. Ultra-thin sections were cut with a diamond knife using a Leica UC7 ultramicrotome at a thickness of 50 nm and collected on slotted grids coated with formvar and carbon. The ultra-thin sections were then stained with 3% uranyl acetate and Reynold’s lead citrate prior to imaging. Data was acquired in a Hitachi HD2300 STEM at 200 kV utilizing phase contrast transmission mode and high angle annular dark field (HAADF).

### Transmission electron microscopy

Samples were prepared and imaged via negative-stain TEM as described in-depth previously [11]. In brief, purified MCP samples were fixed on 400 mesh Formvar-coated copper grids (EMS Cat# FF400-Cu) using 2% (v/v) glutaraldehyde. Fixed samples were washed in deionized water and stained using 1% (w/v) uranyl acetate. Samples were dried and stored until imaging. Fixed and stained MCPs were imaged with the Hitachi HT-7700 Biological S/TEM Microscope and Galtan Orius 4k x 2.67k digital camera at the Northwestern Electron Probe Instrumentation Center (EPIC).

### SDS-PAGE and western blot

For SDS-PAGE separation of purified MCPs, samples were normalized based on total protein concentration. For strains with low yields, the maximum amount of protein possible was loaded on the gel. Samples were boiled at 95 °C in Laemmli buffer for 5 minutes and loaded onto a 15% Tris-glycine mini gels. The gels were run at 120 V for 90 minutes and then stained with Coomassie Brilliant Blue R-250.

Western blots were performed on OD_600_-normalized samples of whole cell lysate. A 1 mL sample of induced culture was pelleted and stored at −20 °C until use. Pellets were resuspended in 25 mM Tris base, 192 mM glycine, 1.1% (w/v) SDS to an OD_600_ of 3.0. Samples were then mixed with 4X Laemmli buffer at 3:1 volume ratio and boiled for 5 minutes at 95 °C. Boiled samples were run on a 15% Tris-glycine gel at 130 V for 80 minutes and transferred to a PVDF membrane at 90 V for 11 minutes by wet transfer with Bio-Rad Mini Trans-Blot Cell. Membranes were blocked in TBS-T (20 mM Tris (pH 7.5), 150 mM sodium chloride, 0.05% (v/v) Tween 20)) with 5% (w/v) dry milk for 1 hour at room temperature. Blocked membranes were incubated overnight at 4 °C with a 1:6666 dilution of mouse anti-FLAG primary antibody (MilliporeSigma Cat# F3165) in TBS-T with 5% (w/v) dry milk. Membranes were washed with TBS-T and incubated in a 1:1000 dilution of HRP-conjugated goat anti-mouse secondary antibody (Invitrogen Cat# 32430) in TBS-T for two hours at room temperature. Finally, membranes were developed using SuperSignal^TM^ West Pico PLUS Chemiluminescent Substrate (Thermo Fisher Scientific) and imaged using the Bio-Rad ChemiDoc XRS+ System.

## Abbreviations

MCP: Bacterial microcompartments
BMC: bacterial microcompartment
BMV: bacterial microcompartment vertex
BMC-H: bacterial microcompartment hexamer
BMC-P: bacterial microcompartment pentamer
BMC-T: bacterial microcompartment trimer
1,2-PD: 1,2-propanediol
Pdu: 1,2-propanediol utilization
ssD: signal sequence from PduD
ssP: signal sequence from PduP
TEM: transmission electron microscopy

## Acknowledgements

The authors would like to thank members of the Tullman-Ercek group for insightful discussions and advice during the planning and preparation of this manuscript. Specifically, we would like to thank Dr. Lisa A. Burdette, Dr. Carolyn E. Mills, and Taylor M. Nichols for helpful feedback on the manuscript composition. We would also like to thank Eric Roth for his expertise in carrying out the ultra-thin section TEM experiments. This work was supported by the National Science Foundation (award MCB1150567 to DTE), the Army Research Office (grant W911NF-19-1-0298 to DTE), and the Department of Energy (grant DE-SC0019337 to DTE). NWK was supported by the National Science Foundation Graduate Research Fellowship Program (grant DGE-1842165), and by the National Institutes of Health Training Grant (T32GM008449) through Northwestern University’s Biotechnology Training Program. HWR was funded in part by Summer Undergraduate Research Grant though the Office of Undergraduate Research at Northwestern University (grant 586SUMMER1812972). This work made use of the EPIC facility of Northwestern University’s NUANCE Center, which has received support from the Soft and Hybrid Nanotechnology Experimental (SHyNE) Resource (NSF ECCS-1542205); the MRSEC program (NSF DMR-1720139) at the Materials Research Center; the International Institute for Nanotechnology (IIN); the Keck Foundation; and the State of Illinois, through the IIN.

## Author Contributions

**NWK**: Conceptualization, methodology, validation, formal analysis, investigation, resources, writing – original draft, writing – review & editing, visualization, supervision, project administration, funding acquisition. **SPI**: Conceptualization, methodology, validation, formal analysis, investigation, resources, writing – original draft, writing – review & editing, visualization, project administration. **MSL**: Conceptualization, methodology, investigation, resources, writing-review & editing. **HWR**: Conceptualization, methodology, formal analysis, investigation, writing – review & editing. **DTE**: Conceptualization, validation, resources, writing – original draft, writing – review & editing, supervision, project administration, funding acquisition.

