## Supplemental Information for "Self-assembling shell proteins PduA and PduJ have essential and redundant roles in bacterial microcompartment assembly"

### Supplemental Figures

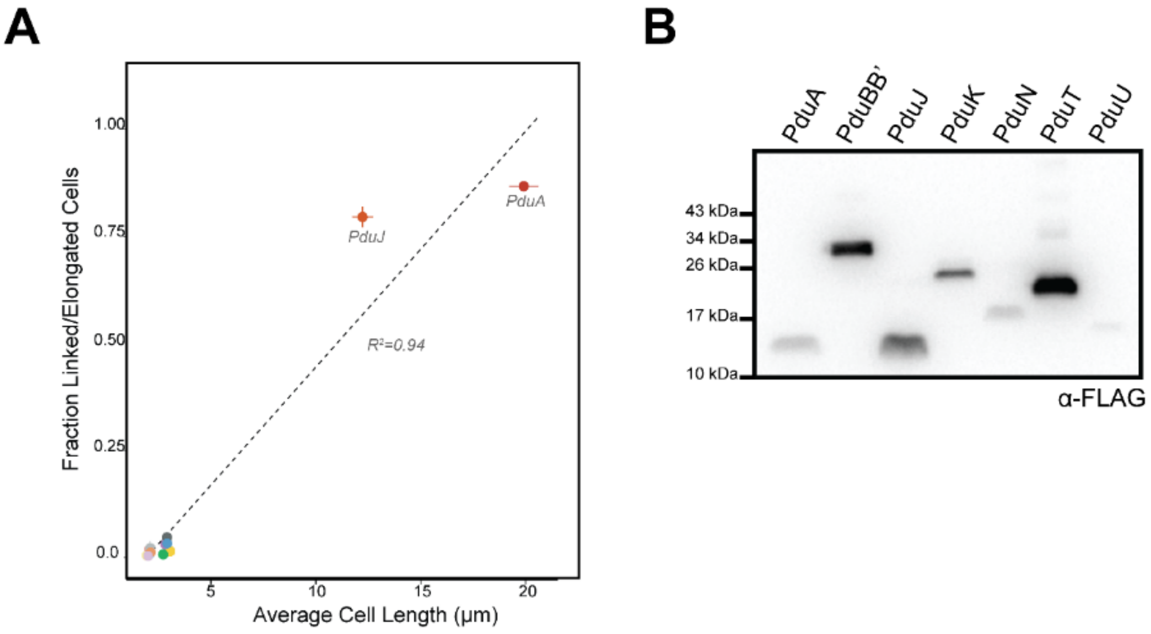


**Figure S1 – Expression and quantification of Pdu shell protein assembly.** (A) Correlation of average cell length (μm) and fraction of linked or elongated cells showing positively correlation of $R^2$=0.94. (B) Western blot of whole cell lysate from cells overexpressing FLAG-tagged Pdu shell proteins. Lanes corresponding to each shell protein are shown across the top of the blot. Samples were normalized by cell density to an $OD_{600}$ of 3.0.

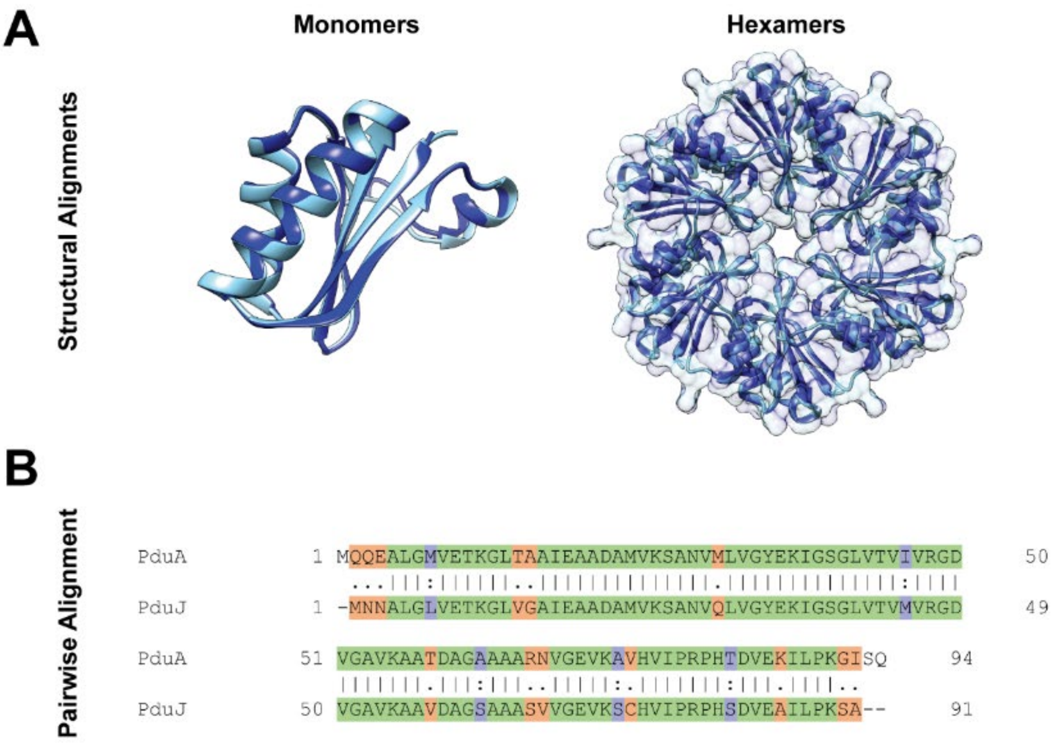


**Figure S2 – Alignments of PduA and PduJ.** (A) Structural alignment of PduA/PduJ monomers (left) and hexamers (right). PduA PDB ID: **3NGK** [1]. PduJ PDB ID: **5D6V** [2]. Structures were visualized using Chimera [3]. (B) Pairwise alignment of PduA and PduJ proteins from *S. enterica* is shown below. The sequences are 77.7% (73/94) identical with a Root Mean Square Deviation of 0.292. [4] Green = identical residues; purple = highly similar residues; orange = weakly similar residues.

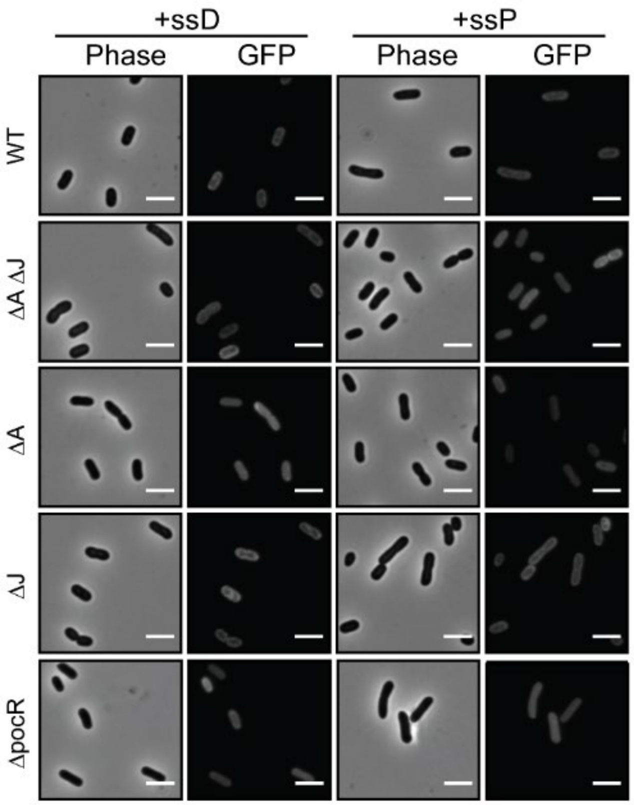


**Figure S3 – Fluorescence microscopy in strains with uninduced MCPs.** Phase contrast and fluorescence microscopy images from WT, ΔA, ΔJ, ΔA ΔJ, and ΔpocR strains expressing either ssD-GFP or ssP-GFP but uninduced for MCP formation (no 1,2-PD added). Scale bars = 3 μm.

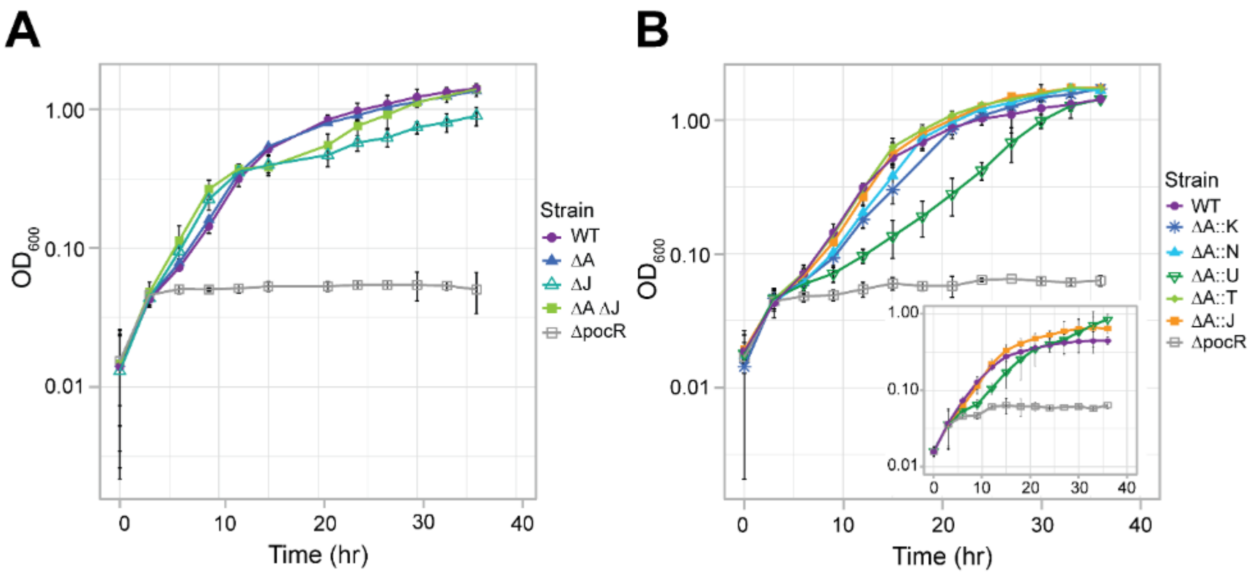


**Figure S4 – PduA and PduJ deletion and substitutions have varied effects on growth.** (A) Growth curve of WT, ΔA, ΔJ, ΔA ΔJ, and ΔpocR strains in NCE media with 55 mM 1,2-PD as a carbon source and supplemented with 150 nM adenosylcobalamin (AdoB$_{12}$). Measurements from three biological replicates were taken and error bars represent 95% confidence interval. (B) Growth curve of WT, ΔA::X substitutions, and ΔpocR strains in NCE media with 55 mM 1,2-PD as a carbon source and supplemented with 150 nM AdoB$_{12}$. (B, Insert) Growth curve of WT, ΔA::U, ΔA::J, ΔpocR strains in NCE media with 55 mM 1,2-PD and 20 nM AdoB$_{12}$. Measurements from at least three biological replicates were taken and error bars represent 95% confidence interval.

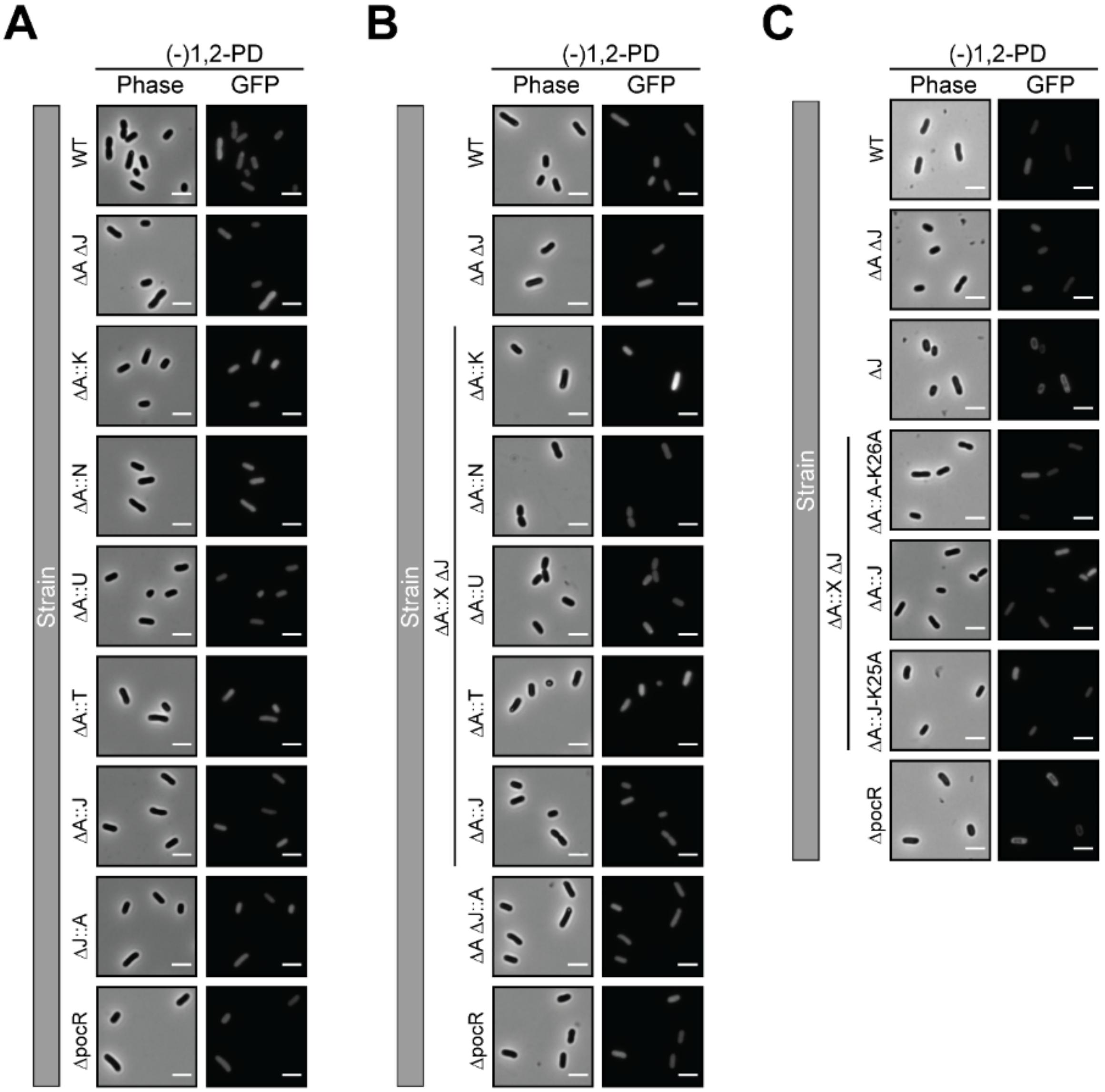


**Figure S5 – Only diffuse ssD-GFP observed in single and double knockout substitution strains when MCP formation uninduced.** Phase contrast and GFP fluorescence microscopy images of various strains expressing ssD-GFP but MCP formation not induced ((-) 1,2-PD). (A) WT, ΔA ΔJ, ΔA::X (X = K,N,U,T, or J), ΔJ::A, and ΔpocR strains. (B) WT, ΔA ΔJ, ΔA::X ΔJ (X = K,N,U,T, or J), ΔA ΔpduJ::A, and ΔpocR strains. (C) WT, ΔA ΔJ, ΔJ, ΔA::A-K26A ΔJ, ΔA::J ΔJ, ΔpduA::J-K25A ΔJ, and ΔpocR strains. Row labels indicate the strain and column labels indicate the type of microscopy. Scale bars = 3 μm.

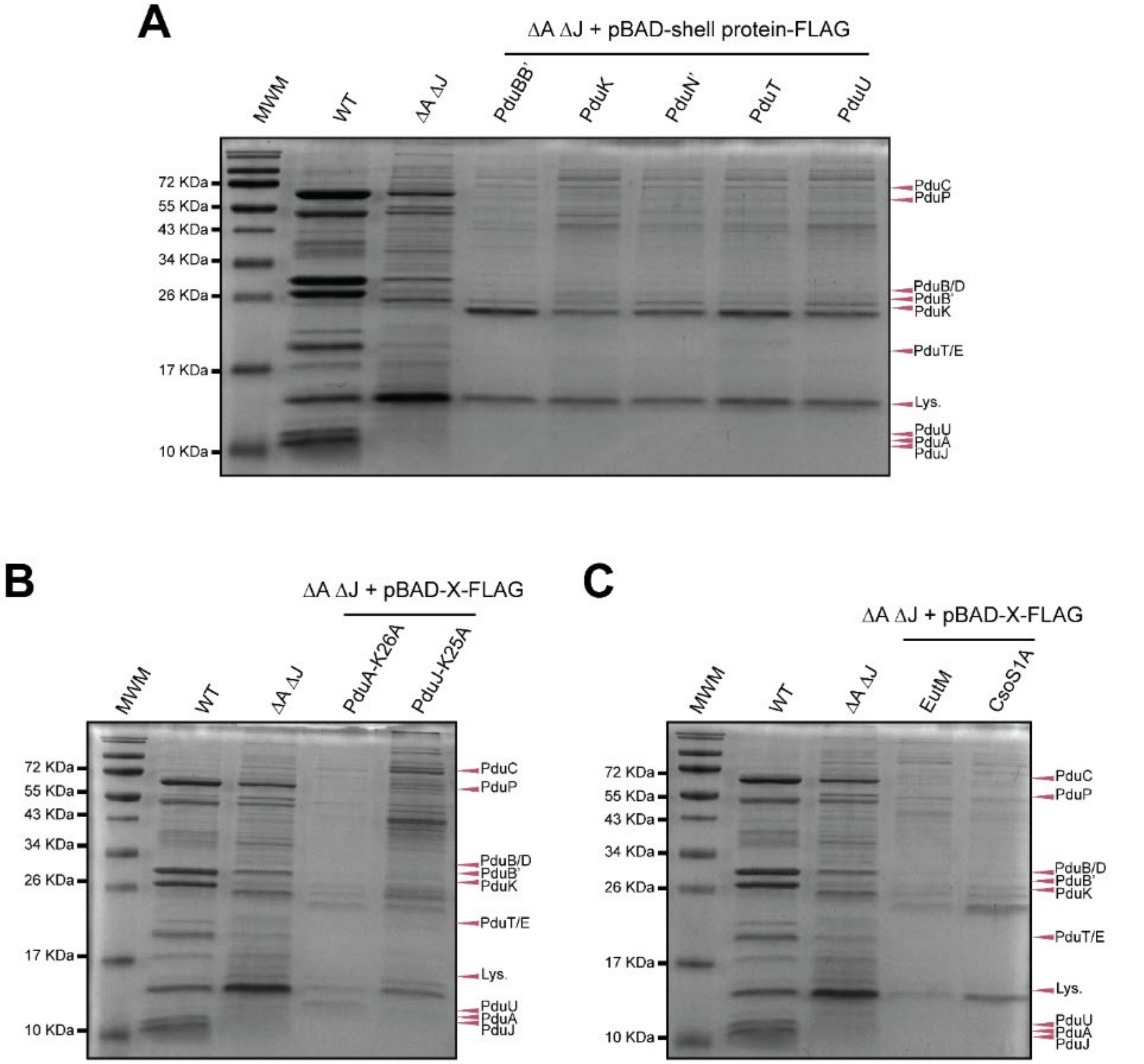


**Figure S6 – Overexpression of other proteins from a plasmid does not rescue MCP formation.** Coomassie-stained SDS-PAGE gel of purified MCPs from different strains with Pdu MCP components labelled. “MWM” is the molecular weight marker and “Lys.” is lysozyme from the lysis buffer. (A) “pBAD + shell protein” indicates that the shell protein indicated on the gel lane was overexpressed in these samples. (B) “pBAD + X” indicates that the assembly-deficient shell protein  labeled on the gel lane was overexpressed in these samples. “MWM” is the molecular weight marker and “Lys.” is lysozyme from the lysis buffer. (C) “pBAD + X” indicates that the PduA homolog indicated on the gel lane was overexpressed in these samples.

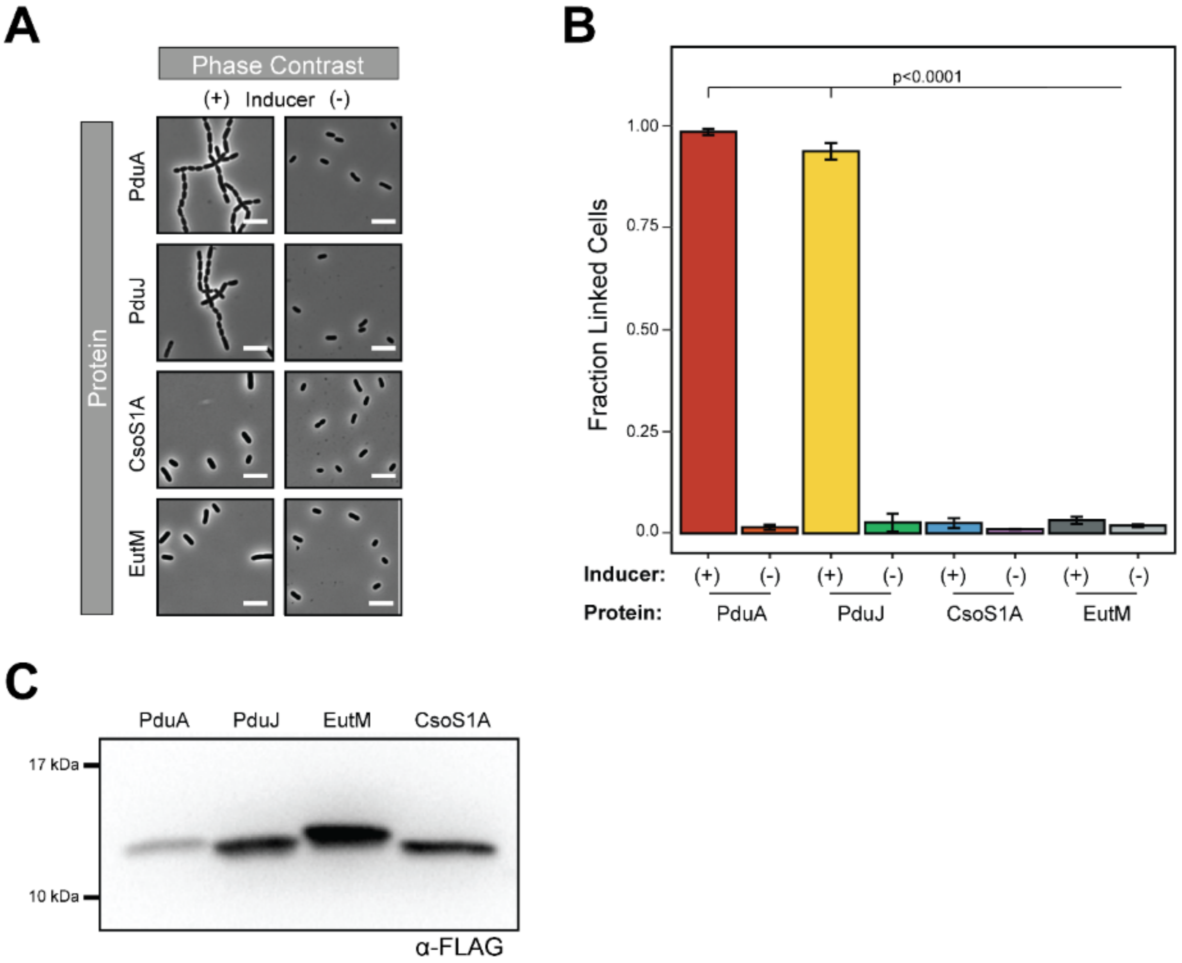


**Figure S7 – PduA homologs EutM and CsoS1A do not self-assemble in the same manner as PduA and PduJ.** (A) Phase contrast microscopy images of cells overexpressing PduA, PduJ, EutM, and CsoS1A. (B) Fraction of linked or elongated cells in population of cells induced (+) or uninduced (-) for overexpression of PduA, PduJ, EutM, or CsoS1A. Cells overexpressing (+ inducer) PduA or PduJ have a significantly greater fraction of cells in linkages of three or more cells or longer than 6 μm ($p < 0.001$, t-test). Counts were taken from three biological replicates (>60 cells counted per strain per replicate), and error bars represent standard error of the means. (D) Western blot of whole-cell lysate from cells overexpressing FLAG-tagged PduA, PduJ, EutM, and CsoS1A.

**Table S1.** Doubling time ($t_d$) of single knockout substitution strains in NCE with 55 mM 1,2-PD and 150 nM $AdoB_{12}$.

| **Strain** | **$t_d$ (hr)** | **p-value[a]** |
|---|---|---|
| WT | 3.22 ± 0.23 | |
| *ΔA::K* | 4.34 ± 0.33 | 0.00017 |
| *ΔA::N* | 3.80 ± 0.36 | 0.007 |
| *ΔA::U* | 5.49 ± 0.09 | < 0.0001 |
| *ΔA::T* | 3.13 ± 0.28 | 0.45 |
| *ΔA::J* | 3.27 ± 0.36 | 0.72 |

[a]With respect to WT. Error is 95% confidence interval of the mean

**Table S2.** Plasmids used in this study.

| **Name** | **Plasmid** | **Origin** | **Resistance** |
|---|---|---|---|
| pSIM6 [5] | λ Red system repressed by cI857 | pSC101 *repA[ts]* | Ampicillin |
| CMJ069 | pBAD33t-ssD-GFPmut2 | p15A | Chloramphenicol |
| EYK208 | pBAD33t-ssP-GFPmut2 | p15A | Chloramphenicol |
| CMJ138 | pBAD33t-PduA-FLAG | p15A | Chloramphenicol |
| CMJ140 | pBAD33t-PduBB'-FLAG | p15A | Chloramphenicol |
| CMJ141 | pBAD33t-PduJ-FLAG | p15A | Chloramphenicol |
| CMJ142 | pBAD33t-PduK-FLAG | p15A | Chloramphenicol |
| CMJ144 | pBAD33t-PduN-FLAG | p15A | Chloramphenicol |
| CMJ145 | pBAD33t-PduT-FLAG | p15A | Chloramphenicol |
| CMJ146 | pBAD33t-PduU-FLAG | p15A | Chloramphenicol |
| NWKp016 | pBAD33t-PduA-K26A-FLAG | p15A | Chloramphenicol |
| NWKp005 | pBAD33t-PduJ-K25A-FLAG | p15A | Chloramphenicol |
| NWKp021 | pBAD33t-EutM-FLAG | p15A | Chloramphenicol |
| NWKp022 | pBAD33t-CsoS1A-FLAG | p15A | Chloramphenicol |

**Table S3.** Primers used in this study.

| Name | Purpose | Description | Sequence* |
|---|---|---|---|
| MFSP 105 | Recombineering | Amplify *cat/sacB* with homology upstream to *pduA* For | ttatagtcccaactatcggaacactccatgcgaggtctttTGTGACGGAAGATCACTTCG |
| MFSP 106 | Recombineering | Amplify *cat/sacB* with homology at C-terminus of *pduA* Rev | ttcccttcggtaagatttttctacatcggtgtgagggcgATCAAAGGGAAAACTGTCCATAT |
| NWKo 328 | Recombineering | Amplify *cat/sacB* with homology to *pduJ* For | atgaataacgcactgggactggttgaaacaaaagggctggTGTGACGGAAGATCACTTCG |
| SPIP 055 | Recombineering | Amplify *cat/sacB* with homology upstream to *pduJ* For | ctttcgggatctccatgcttaatcacaggagaacggcagtTGTGACGGAAGATCACTTCG |
| NWKo 329 | Recombineering | Amplify *cat/sacB* with homology at C-terminus of *pduA* Rev | ggctgatttcggtaaaatggcctcaacatcgctgtgcATCAAAGGGAAAACTGTCCATAT |
| MFSP 333 | Recombineering | *ΔpduA* knockout cat/sacB For | ttcccttcggtaagatttttctacatcggtgtgagggcgaaagacctcgcatggagtgttc |
| MFSP 334 | Recombineering | *ΔpduA* knockout cat/sacB Rev | ttatagtcccaactatcggaacactccatgcgaggtctttcgccctcacaccgatgtag |
| NWKo 334 | Recombineering | *ΔpduJ* KO cat/sacB For | gaataacgcactgggactggttgaaacaaagggctgggcacagcgatgttgaggccattttaccgaaatcagcc |
| SPIP 053 | Recombineering | Amplify *pduK* with homology at *pduA* For | ttatagtcccaactatcggaacactccatgcgaggtctttATGGCGAATAAGGAGCACCG |
| SPIP 054 | Recombineering | Amplify *pduK* with homology at *pduA* Rev | ttcccttcggtaagatttttctacatcggtgtgagggcgTTACGCTTCACCTCGCTTGC |
| MFSP 368 | Recombineering | Amplify *pduN* with homology at *pduA* For | ttatagtcccaactatcggaacactccatgcgaggtctttATGCATCTGGCACGAGTCAC |
| MFSP 369 | Recombineering | Amplify *pduN* with homology at *pduA* Rev | ttcccttcggtaagatttttctacatcggtgtgagggcgTTAACACGAAAGCGTATCTACAATGCC |
| MFSP 370 | Recombineering | Amplify *pduU* with homology at *pduA* For | ttatagtcccaactatcggaacactccatgcgaggtctttATGGAAAGACAACCGACAACGG |
| MFSP 371 | Recombineering | Amplify *pduU* with homology at *pduA* Rev | ttcccttcggtaagatttttctacatcggtgtgagggcgTTACGTCCGGGTGATCGAGC |
| MFSP 372 | Recombineering | Amplify *pduT* with homology at *pduA* For | ttatagtcccaactatcggaacactccatgcgaggtctttATGTCTCAGGCTATAGGAATTTTAGAACTCAC |
| MFSP 373 | Recombineering | Amplify *pduT* with homology at *pduA* Rev | ttcccttcggtaagatttttctacatcggtgtgagggcgTTACCCCTCCACCATCTGTCG |

| | | | |
|---|---|---|---|
| MFSP 374 | Recombineering | Amplify *pduJ* with homology at *pduA* For | ttatagtcccaactatcggaacactccatgcgaggtctttATGAATAACGCACTGGGACTGG |
| MFSP 375 | Recombineering | Amplify *pduJ* with homology at *pduA* Rev | ttcccttcggtaagatttttctacatcggtgtgagggcgTTAGGCTGATTTCGGTAAAATGGCC |
| SPIP 056 | Recombineering | Amplify *pduA* with homology at *pduJ* For | ctttcgggatctccatgcttaatcacaggagaacggcagtATGCAACAAGAAGCACTAGGAATGG |
| SPIP 057 | Recombineering | Amplify *pduA* with homology at *pduJ* Rev | ggctgatttcggtaaaatggcctcaacatcgctgtgcTCATTGGCTAATTCCCTTCGGTA |
| SPIP 058 | Recombineering | Amplify *eutM* with homology at *pduA* For | ttatagtcccaactatcggaacactccatgcgaggtctttATGGAAGCATTAGGAATGATTGAAACC |
| SPIP 059 | Recombineering | Amplify *eutM* with homology at *pduA* Rev | ttcccttcggtaagatttttctacatcggtgtgagggcgTCAAATGTTGCTGTCGCCTT |
| NWKo 418 | Recombineering | Amplify pduA with homology at pduA For | gcatcttcttatagtcccaactatcggaacactccatgcgaggtctttatgcaacaagaagcactaggaatg |
| NWKo 419 | Recombineering | Amplify pduA with homology at pduA Rev | gggcaatcacctgcgccatgatctgttccaccagctcattgctgctcattggctaattcccttcggtaaga |
| NWKo 466 | Recombineering/ sequencing | Amplify from upstream of pduA locus For | CTGCGAACCTGTCTCC |
| NWKo 467 | Recombineering/ sequencing | Amplify from downstream of pduA locus Rev | CGTCTCTCGTATAGGTTGG |
| NWKo 017 | Site-directed mutagenesis | PduA-K26A mutagenesis For | CCGCTGATGCAATGGTTGCGTCAGCCAATGTGATG |
| NWKo 018 | Site-directed mutagenesis | PduA-K26A mutagenesis Rev | CATCACATTGGCTGACGCAACCATTGCATCAGCGG |
| NWKo 025 | Site-directed mutagenesis | PduJ-K25A mutagenesis For | CGCCGATGCAATGGTTGCGTCCGCCAACGTACAGC |
| NWKo 026 | Site-directed mutagenesis | PduJ-K25A mutagenesis Rev | GCTGTACGTTGGCGGACGCAACCATTGCATCGGCG |
| NWKo 330 | Sequencing | Amplify from upstream of *pduJ* | GATCGACACTCGCTGGTCGTGCAT |
| NWKo 331 | Sequencing | Amplify from downstream of *pduJ* | GGCCACATCACCGGTAATTTTAATCACCAT |

*Homology region is underlined
